# Targeted modulation of IGFBP5/IGF1, THPO, and P38 MAPK signaling are potent therapeutic strategies generalizable for mitochondrial respiratory chain disease and osteosarcoma

**DOI:** 10.64898/2026.07.20.739625

**Authors:** Kelsey Keith, Min Peng, Cristina Remes, Victoria Miranda, Nicholas Wachowski, Melis Kose, Shrikant Dharaskar, Suraiya Haroon, Agustin B. Velasco, Priya Sivaramakrishnan, Donna Iadarola, Sundeep Dugar, Marni J. Falk

## Abstract

Primary mitochondrial diseases (PMD) have limited disease-modifying therapies, currently applicable to only 3 of over 400 discrete gene disorders. Cycloheximide (CHX) is a global cytosolic translation inhibitor we previously reported to rescue PMD preclinical models, although its toxicity precluded clinical development. To identify specific mediators underlying CHX treatment benefit in PMD, SOMAscan-based proteomics was performed in complex I deficient and genetic disease fibroblast cell line models grown in galactose. Thrombopoietin (THPO) and insulin-like growth factor binding protein 5 (IGFBP5) were the only two differentially regulated proteins, together with ERK/MAPK pathway dysregulation, identified upon CHX treatment in PMD versus healthy control cells. THPO inhibition by siRNA or pharmacologic approaches rescued stress-induced viability loss in patient fibroblasts having diverse PMD gene etiologies, and significantly improved mitochondrial stress, linear growth, and neuromuscular function in a classical *ndufs2^-/-^ C. elegans* model. IGFBP5 overexpression by lentiviral or mRNA approaches rescued cell viability across distinct PMD gene etiologies, as did IGF1 pharmacologic inhibition across both PMD mutant and *C. elegans* models. MAPK pharmacologic inhibition rescued multiple distinct complex I disease cells’ survival, as well as mitochondrial stress in *SLC25A46^-/-^ C. elegans*. Combination therapies targeting multiple of these glucose signaling pathway proteins, together with glucose and N-acetylcysteine, yielded superior therapeutic benefit in complex I disease cell and *C. elegans* models. Additionally, single or combined pharmacologic inhibition of THPO or IGF1 significantly enhanced primary and metastatic osteosarcoma cell death. Collectively, targeted small molecule and genetic modulation of THPO, IGF1, or MAPK recapitulated the significant therapeutic benefit of CHX in PMD, while avoiding global translation inhibition. These novel PMD therapies likely confer benefit by attenuating MAPK-driven autophagy and potentially promoting noncanonical glucose uptake, improving cellular energy balance. Overall, these glucose signaling cellular pathway targets hold broad therapeutic promise for PMD patients, warranting further clinical research development.

## INTRODUCTION

Among the 1,136 genes that encode proteins or RNAs that have been experimentally verified to localize within mitochondria [1], nearly all are encoded by the nuclear genome except for 37 mitochondrial DNA (mtDNA) genes that yield 13 protein-coding messenger RNAs and 24 non-coding transfer and ribosomal RNAs [2]. Primary mitochondrial diseases (PMD), which collectively have a minimal population prevalence of 1 in 4,300 individuals across all ages and backgrounds [3], are now recognized to occur by any mode of inheritance from pathogenic variants in over 400 genes [4]. PMD demonstrates extreme clinical heterogeneity, potentially affects all body systems, and has variable penetrance across a given gene’s variants, whether it be a nuclear or mtDNA gene disorder. mtDNA variant related clinical variability is partially explained by heteroplasmy, reflecting the mitochondrial genome percentage carrying a specific mutation in a given tissue [5]. Recently, the first disease modifying treatments for mitochondrial disease have received regulatory approval by the Food and Drug Administration (FDA) for 3 single nuclear gene disorders, including Raloxone (omevaloxalone) for Friedrich’s ataxia due to *Frataxin* mutations; Forzinity (elamipretide) for Barth syndrome due to impaired cardiolipin biosynthesis [6]; and Kygevvi (doxecitine and doxribtimine combination nucleosides) for thymidine kinase 2 deficiency (TK2d) due to mutation in the TK2 nuclear gene [7]. However, no US-FDA approved treatments exist for the general clinical indication of PMD. Rather than targeting the root drivers of disease, current standards of care for most gene etiologies and clinical manifestations of PMD relate to symptomatic management and using empiric therapies to enhance cellular stress resilience and restore intermediary metabolite deficiencies by targeting common downstream aspects of cellular pathophysiology such as increased oxidative stress, NADH/NAD^+^ redox imbalance, one carbon metabolism deficiency, and nitric oxide dysregulation [8–11]. Indeed, a major focus of clinical therapeutic development for PMD has attempted to repurpose existing therapies and/or develop therapies that target oxidative stress, NAD^+^ metabolism, or nutrient-sensing signaling pathways such as PPAR-delta involved in regulating intermediary metabolism [11–18].

Cytosolic protein translation in PMD was consistently recognized by our group previously to be significantly dysregulated [19–21]. To this end, our research group discovered that low-micromolar concentrations of cycloheximide (CHX), as well as actinomycin and anisomycin, effectively rescued viability, proteotoxic stress, respiratory capacity, and inversely dysregulated mitochondrial translation and biogenesis across diverse pharmacologic and genetic cellular models of PMD [21]. However, CHX inhibition of protein translation involves direct ribosome binding [22], with global changes in cellular regulation that appeared unnecessary for its therapeutic effect but likely contributed to its known adverse side effects as a chemotherapeutic agent. Transcriptome profiling and early mass-spectrometry-based proteomics analyses failed to reveal its specific mechanism, with the specific identity remaining elusive of several proteins that paradoxically appeared to be upregulated on ^35^S-methionine translation activity analyses in PMD cells treated with CHX [21, 23].

In an effort to identify novel therapeutic targets underlying CHX’s unexpected therapeutic benefit in PMD, we sought to harness advanced proteomics technology to unmask the identity of specific proteins that are dysregulated in human PMD patient cells and rescued by CHX treatment. Specifically, we evaluated a SomaLogic DNA aptamer proteomics assay that had been originally developed for blood profiling in a new matrix of primary human fibroblast cells. Bioinformatics analyses were performed to evaluate the identity of specific proteins showing significant differential regulation by either a mitochondrial respiratory chain complex I inhibitor (rotenone), or by nuclear gene-based complex I diseases, with significant reversal upon low-micromolar concentration CHX treatment. Surprisingly, only two candidate proteins emerged from these proteomics-wide analyses as therapeutic targets for mitochondrial complex I disease, with signaling pathway level analyses identifying an additional mechanistic target. These three protein targets were validated by in vitro gene overexpression and/or inhibition studies in diverse cellular PMD models, as well as in a *C. elegans gas-1(fc21)* model of *NDUFS2^-/-^* complex I disease. Given the known therapeutic role of CHX as a chemotherapy, these three therapeutic targets were also evaluated in human cell models of osteosarcoma.

## RESULTS

PMD patient (n=3, including *NDUFS1^-/-^*, *NDUFS4^-/-^*, and *NDUFS8^-/-^*) and healthy control fibroblasts (n=3 asymptomatic parents of PMD patients, as detailed in **Supp. Fig 1A**) were cultured in standard media alone or co-treated with rotenone, a mitochondrial respiratory chain complex I inhibitor that chemically mimics mitochondrial disease. All cell lines were initially grown in a glucose-rich media (5.6 mM) and then switched to galactose (10 mM). Galactose media unmasks mitochondrial respiratory chain deficiencies by forcing conversion of galactose to glucose, thereby lowering glucose availability for glycolysis and forcing the cultured cells to rely to a greater extent on mitochondrial oxidative phosphorylation (OXPHOS) for ATP generation. Indeed, PMD cells compensate for their inherent OXPHOS dysfunction by relying on glycolysis for energy production, as glucose itself is therapeutic in PMD models [24–26].

**Fig 1.**
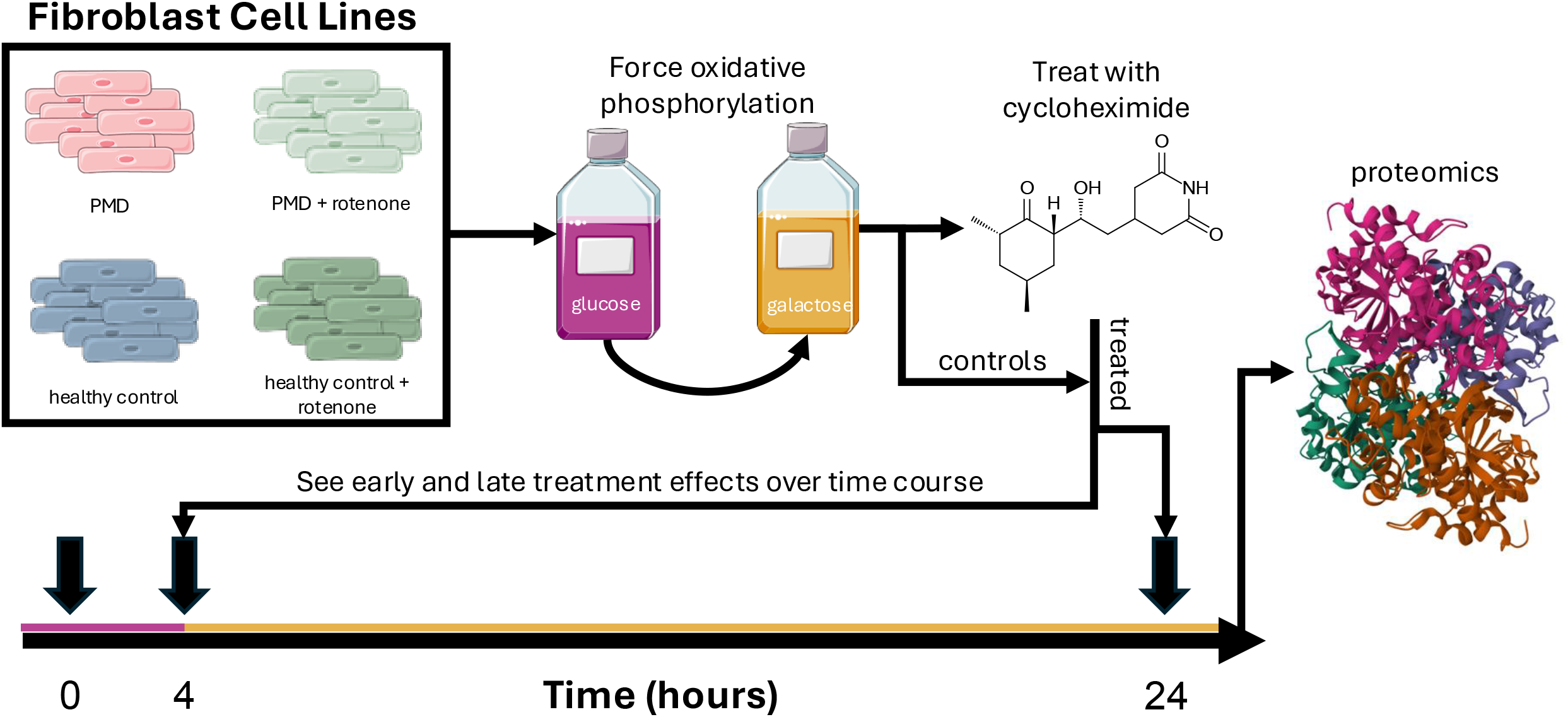
Proteomics Experimental Design Schematic. Primary human fibroblast cell lines from mitochondrial disease (PMD) patients with mitochondrial respiratory chain complex I subunit mutations (n=3) and healthy parental controls (conveyed by “P1”, n=3) were studied. All cells were initially cultured in glucose-rich media and then changed to growth in galactose-only media for up to 24 hours to force reliance on mitochondrial OXPHOS capacity for energy generation. To exacerbate complex I dysfunction, cell lines were treated with rotenone, which is a potent complex I specific chemical inhibitor. Cycloheximide was co-incubated in galactose media, as a chemical inhibitor of cytosolic protein translation. Glucose media exposed samples were collected at baseline at 0 hours, as glucose masks PMD effects. All other treatments were collected in galactose media at 4 and 24 hrs. Protein was extracted from all cell lines and sent to SomaLogic for aptamer-based proteomics profiling.

Simultaneous with the switch to galactose media, cells were treated with cycloheximide (CHX). All conditions were sampled at 4-hours and 24-hours post-media switch and then sent to SomaLogic for proteomics (**Fig. 1**).

As fibroblast cell lines presented a new matrix for SomaLogic, modification was made to their proprietary DNA aptamer proteomics assay to optimize for the dynamic range of this new sample type. All samples passed quality control analysis (Supp. Fig 1B). However, samples from *NDUFS8^-/-^* (mitochondrial complex I disease patient) and *NDUFS8^+/-^* (healthy parent carrier) cell lines were initially collected and proteomics analysis was performed, but these two samples were subsequently recognized to lack one of the disease-causing alleles and therefore removed from downstream data analysis as genetic mutation confirmation was not possible to perform post-hoc and PCA analysis showed a clustering effect (**Supp. Fig. 1A**). Differential proteomics analysis was carried out following SomaLogic recommendations. For healthy and disease groups at both 4-hours and 24-hours, comparative proteomic results in each treatment condition were performed relative to untreated cells in galactose media, as follows: galactose media vs glucose media, rotenone treatment vs galactose media, CHX treatment vs galactose media, and combination CHX and rotenone treatment vs galactose media.

All treatment conditions showed a strong effect on protein expression levels. As expected, CHX decreased most protein levels, consistent with its being a well-established global inhibitor of cytosolic translation (**Fig. 2A**). Upon analysis of PMD patient cells (n=2, namely *NDUFS1^-/-^* and *NDUFS4^-/-^*), the switch from glucose to galactose media induced an early stress response evident at 4 hours that persisted at 24 hours, with altered protein and phosphorous metabolism and negative impact on cell adhesion and cell death regulation (**Fig. 2B-C**). Despite CHX increasing mitochondrial disease cell viability [21], proteomic analysis of the top-level pathways showed similar alterations when comparing galactose and glucose media effects in PMD cells (**Fig. 2B-C**) and the effects of CHX vs glucose media in PMD cells (**Fig. 2D-E**).

**Fig 2.**
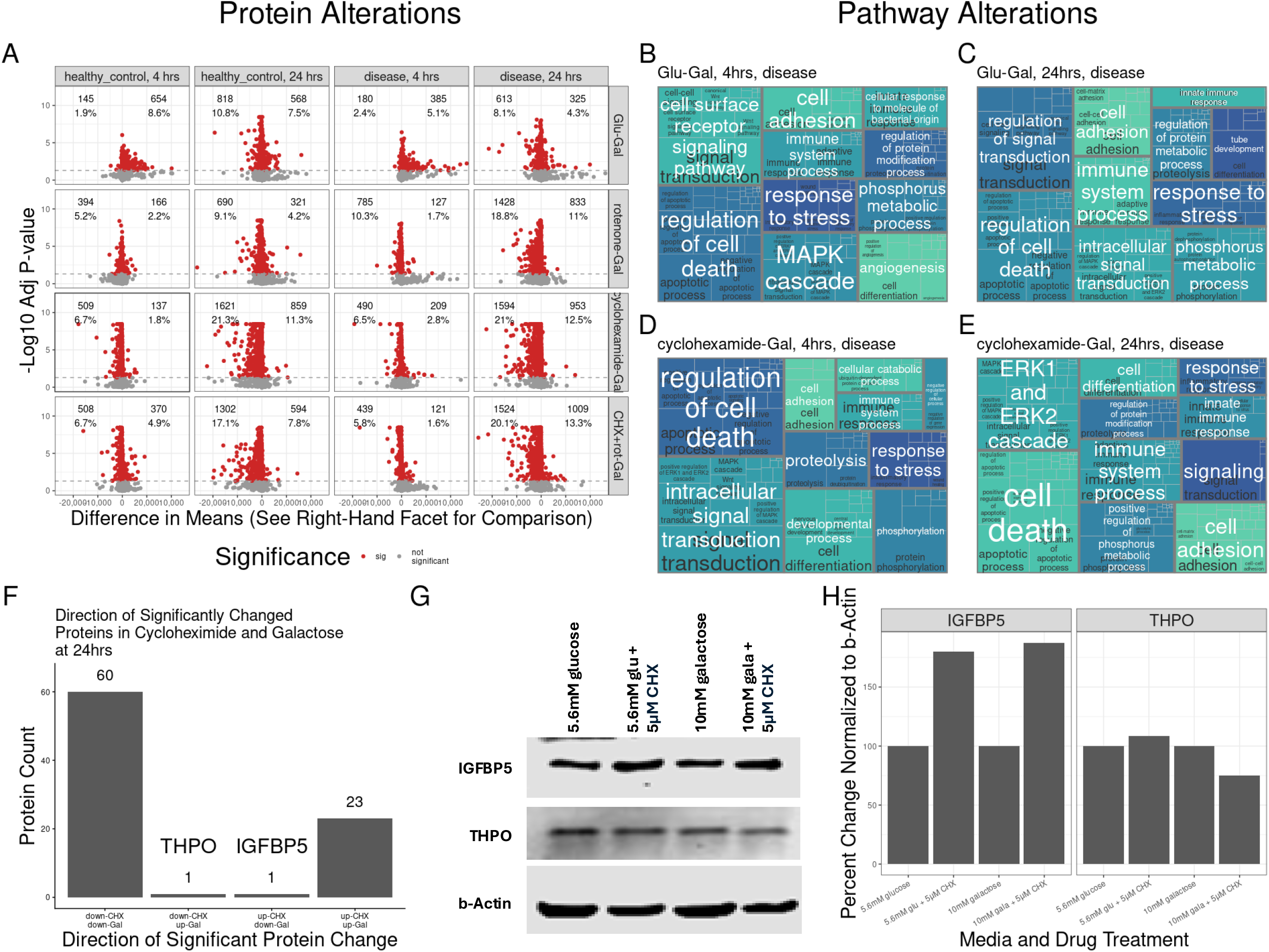
Proteomics Analysis Results. **(A)** Volcano plots display differential protein expression in all conditions tested. Healthy and PMD (‘disease’) fibroblast cell line samples were collected at 4 and 24 hrs. Differences in protein expression between glucose media (GLU), rotenone (ROT), cycloheximide (CHX), and combination rotenone and CHX were determined relative to galactose media (GAL) only as the common reference group. Differential testing was conducted using ANOVA with post-hoc Tukey test, where p values were corrected for multiple testing using Benjamini-Hotchberg analysis. X-axis conveys the difference in means and Y-axis indicates the –log10 of the corrected p-value. Significantly differentially expressed proteins are delineated in red, with the number and percentage of differentially up- and down-regulated proteins shown in the facet for the respective test conditions. Pathway analysis was performed for PMD fibroblast cell line proteomics results comparing **(B)** glucose vs galactose at 4 hours, **(C)** glucose vs galactose at 24 hours, **(D)** CHX at 4 hours, and **(E)** CHX at 24 hours. Treemap plots convey the top 10 parent pathways, which are printed in white with the subpathways grouped behind and colored by the parent pathway with names printed in black, where possible. **(F)** Bar plot showing the number of significantly differentially expressed proteins and their relative direction of change in PMD patient cells cultured at 24 hours in galactose vs glucose media alone or with CHX treatment Importantly, while most proteins change in the same direction in both conditions (down_down or up_up), only two proteins were differentially expressed between these two conditions (down_up and up_down), namely IGFBP5 (up with CHX) and THPO (down with CHX). **(G)** Western immunoblot analysis validated differential IGFBP5 and THPO expression in complex I deficient *NDUFS4^-/-^* PMD patient fibroblast cells grown in 5.6 mM glucose media, glucose with 5 μM CHX, 10 mM galactose media, and galactose with 5 μM CHX. **(H)** Quantification of western immunoblots shown in Fig 3G performed by ImageJ analysis. Upon CHX treatment, IGFBP5 expression increased in both glucose and galactose media while THPO expression decreased only in galactose media, independently validating the SomaLogic proteomics analysis results.

Interestingly, while MAPK cascade expression in PMD cells grown in galactose media was significantly altered at 4 hours, CHX treatment of PMD cells grown in galactose media only induced significant alteration in the ERK1/2 (subsection of MAPK) cascade at 24 hours. These results suggest that CHX treatment of PMD cells delayed ERK/MAPK pathway activation.

While regulation of cell death, stress response, and metabolism were commonly altered pathways overrepresented in all conditions tested including in healthy cells treated with rotenone (**Supp. Fig. 2**), ERK/MAPK response was conspicuously unique to PMD genetic disease cells (**Fig. 2**).

While CHX globally inhibits cytosolic protein translation, our previous work using ^35^S labeling suggested that some protein levels were surprisingly upregulated after CHX treatment – unfortunately, we were not previously able with existing mass spectrometry based methods to determine the specific identity of those proteins observed on western immunoblot analysis to increase with CHX treatment [21]. Accordingly, we postulated that the therapeutic benefit of CHX identified in PMD cells was a fortunate side effect driven by selective upregulation in a single or small subset of proteins whose expression was prioritized under conditions of global translation inhibition. Therefore, when evaluating 24 hour conditions of galactose vs glucose to CHX vs galactose, we prioritized identification of proteins having significant differential expression between these conditions in PMD cell lines but not in healthy controls. Remarkably, only two such proteins met these stringent criteria: (1) thrombopoietin (THPO), which was upregulated in galactose and downregulated in CHX compared to galactose; and (2) IGFBP5, which was downregulated in glucose and upregulated with CHX treatment compared to galactose (**Fig. 2F**). Western immunoblot analysis in *NDUFS4^-/-^* patient fibroblasts strongly and independently validated the SomaLogic proteomics-based findings (**Fig. 2G-H**), where upon CHX treatment, IGFBP5 expression was increased in both glucose and galactose media while THPO expression decreased only in galactose media. Based on these results, we predicted that targeted knockdown of THPO and/or targeted upregulation of IGFBP5 would recapitulate the therapeutic benefits of CHX in PMD.

THPO is a growth factor whose major role is in platelet production and hematopoiesis but also has other tissue functions. THPO binding induces receptor homodimerization to activate JAK/STAT and MAPK signaling cascades, which subsequently regulate cell proliferation, differentiation and other signaling events [27]. To validate the proteomics findings of THPO playing a potential role as a central therapeutic target in PMD, we used an siRNA knockdown of THPO in sequence-verified PMD disease human patient fibroblast cell lines with pathogenic variants in nuclear-encoded complex I subunit genes, namely *NDUFS8^-/-^*(c.160C>T,c.58G>C) and *NDUFS4^-/-^* (c.377_384del). Under galactose media stress for 72 hours to elicit the unmasked cellular consequences of mitochondrial OXPHOS deficiency, that include reduced cell viability, siRNA-based THPO inhibition significantly rescued cell viability in *NDUFS8^-/-^*fibroblast cells (**Fig. 3A**) and also with the combined stressor of low glutamine (250 µM) in the media in *NDUFS4^-/-^* cells (**Fig. 3B**). Similarly, direct inhibition of THPO in these same two complex I disease fibroblast cell lines using a chemical inhibitor, eltrombopag (ELT) was tested under variable *in vitro* conditions (**Fig. 3C-D**). ELT significantly increased cell viability in *NDUFS8^-/-^* cells when grown in glucose media with higher dose (75 nM) rotenone, as well as in galactose (10 mM) media with lower dose (35 nM) rotenone given its hypersensitivity to complex I inhibitors in galactose (**Fig. 3C**). *NDUFS4^-/-^* cell line viability when grown in galactose plus very low dose rotenone (15 nM) was also significantly improved with 2 µM ELT (**Fig. 3D**).

**Fig 3.**
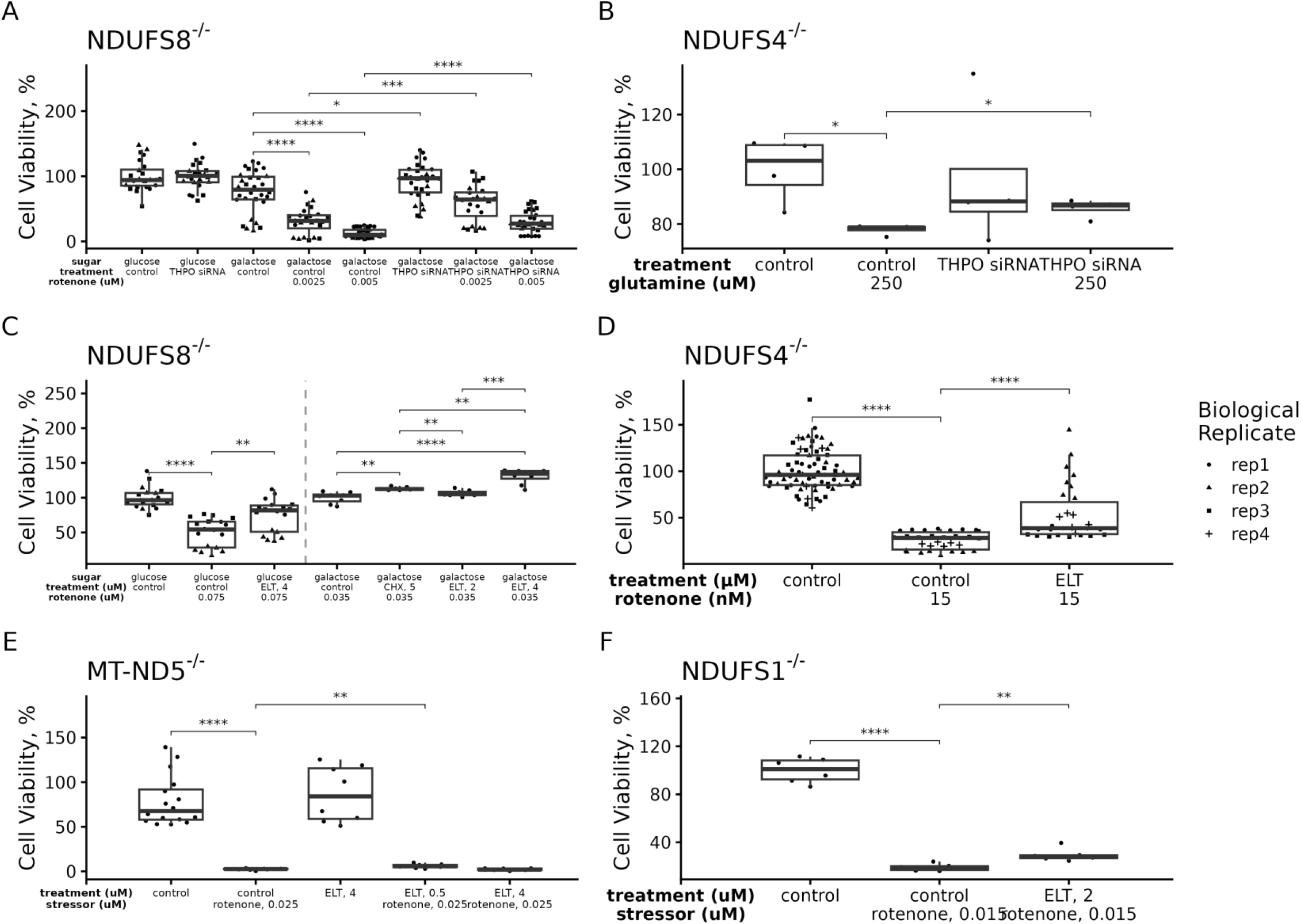
THPO inhibition by siRNA or eltrombopag (ELT) rescued mitochondrial disease cell survival under metabolic stress. **(A)** *NDUFS8^-/-^* fibroblasts cells showed significantly improved cell viability after treatment with THPO siRNA when stressed either in 10 mM galactose media alone or together with either 2.5 nM or 5 nM rotenone. **(B)** *NDUFS4^-/-^* fibroblast cells showed reduced viability in galactose media when culture with low (250 μM) glutamine, which was significantly improved with siRNA inhibition of THPO. **(C)** *NDUFS8^-/-^* fibroblasts cells exposed to high-dose rotenone (75 nM) in glucose or to lower-dose rotenone (35 nM) in galactose had reduced viability, which was significantly increased when treated with a THPO chemical inhibitor, eltrombopag (ELT). ELT (4 µM) treatment effect exceeded that seen upon treatment with 5 µM CHX. **(D)** *NDUFS4^-/-^* cells treated with ELT (2 µM) showed increased survival in galactose media with low-dose rotenone. **(E)** *MT-ND5* fibroblast cells exposed to rotenone (25 nM) had reduced viability that was marginally but significantly treated with ELT (4 µM)**. (F)** *NDUFS1^-/-^* cells grown in galactose with rotenone (15 nM) had reduced viability that was significantly increased with ELT (2 µM) treatment. All panels show results of 72-hour treatment duration, with n=1 biological replicate/condition and 3 technical replicates, unless otherwise specified. Cell viability was tested with a t-test with Benjamini-Hochberg for multiple test correction: *, p < 0.05; **, p < 0.01; ***, p < 0.001; ****, p < 0.0001.

Interestingly, direct THPO inhibition induced a greater increase in cell viability relative to CHX treatment in *NDUFS8^-/-^*cells (**Fig. 3C**). Additional PMD complex I patient fibroblast cell lines were studied harboring pathogenic variants in mitochondrial DNA complex I, starting with subunit *MT-ND5* (m.13513G>A), which was grown in low-dose rotenone (25 nM) to reduce cell viability, but had only a marginal degree of rescue with 0.5 µM ELT (p < 0.01) (**Fig. 3E**).

Similarly, THPO chemical inhibition in the same fibroblasts from the proteomics analysis from a PMD patient with nuclear DNA-encoded complex I subunit *NDUFS1^-/-^* (c.592_594del and c.1727G>A compound heterozygote) grown in galactose had significantly increased but modest improvement in cell viability upon treatment with 2 µM ELT (**Fig. 3F**). Thus, THPO inhibition was validated as a therapeutic target with modest effect in diverse *in vitro* models of complex I-based PMD.

IGFBP5 plays a central role in glucose signaling, where we have identified glucose itself to have therapeutic effects in diverse models of PMD [24–26]. To further validate the potential role of IGFBP5, identified by proteomics analysis, as a central therapeutic target in PMD, IGFBP5 overexpression via lentiviral transfection was performed, demonstrating a significant rescue of cell viability under conditions of metabolic stress in PMD complex I deficient cell lines (**Fig. 4A-B**). Specifically, IGFBP5 overexpression rescued cell viability in *MT-ND5* complex I disease fibroblast cells grown in galactose media stress conditions with high-dose (75 nM) rotenone (**Fig. 4A**). Similarly, IGFBP5 overexpression rescued cell viability in *NDUFS4^-/-^* complex I disease fibroblast cells in galactose media (**Fig. 4B**). Using an independent therapeutic modeling approach, overexpression with IGFBP5 mRNA also significantly rescued *MT-ND5* cell viability both when grown in glucose media with high-dose (75 nM) rotenone, as well as when grown in galactose media with either 5 nM or 7.5 nM rotenone (**Fig. 4C**). Importantly, IGFBP5 acts in the insulin signaling pathway by direct binding and inhibition of IGF1 [28, 29] (**Fig. 4D**). Therefore, we next tested whether a chemical inhibitor of IGF1, OSI-906 (linsitinib), would also rescue PMD cell viability. Indeed, OSI-906 significantly rescued cell viability in PMD disease fibroblast cell lines caused by pathogenic variants in 3 different nuclear-encoded complex I subunit genes (*NDUFS4^-/-^, NUBPL^-/-^* (c.166G>A and c.815-27T>C compound heterozygote)*, NDUFS8^-/-^*) when grown in galactose media with a mutation-specific rotenone concentration (15 nM, 100 nM, 5 nM, respectively) (**Fig. 4E-G**). The same was true for mtDNA-encoded complex I subunit gene, *MT-ND5*, when grown in galactose media (**Fig. 4H**). Beyond complex I disease, OSI-906 (1 µM) treatment also significantly rescued cell viability upon low-dose (2.5 nM) rotenone exposure in galactose media in *FBXL4^-/-^*(c.1067DEL and c.1790A>C compound heterozygous) based mitochondrial disease patient fibroblast cells with multiple respiratory chain complex deficiencies with mtDNA depletion [30, 31], across 3 independent biological replicate experiments (**Fig. 4I**). Similarly, 2 µM OSI-906 significantly rescued cell viability in a *SURF1^-/-^* (c.532_535DEL and c.751C>T compound heterozygous) patient fibroblast line with complex IV deficiency exposed in galactose media to 55 µM sodium azide (**Fig. 4J**). Collectively, these experiments validated both IGFBP5 overexpression and IGF1 inhibition as central therapeutic targets for complex I disease, and extended IGF1 inhibition as a broader therapy for diverse metabolic and genetic etiologies of PMD.

**Fig 4.**
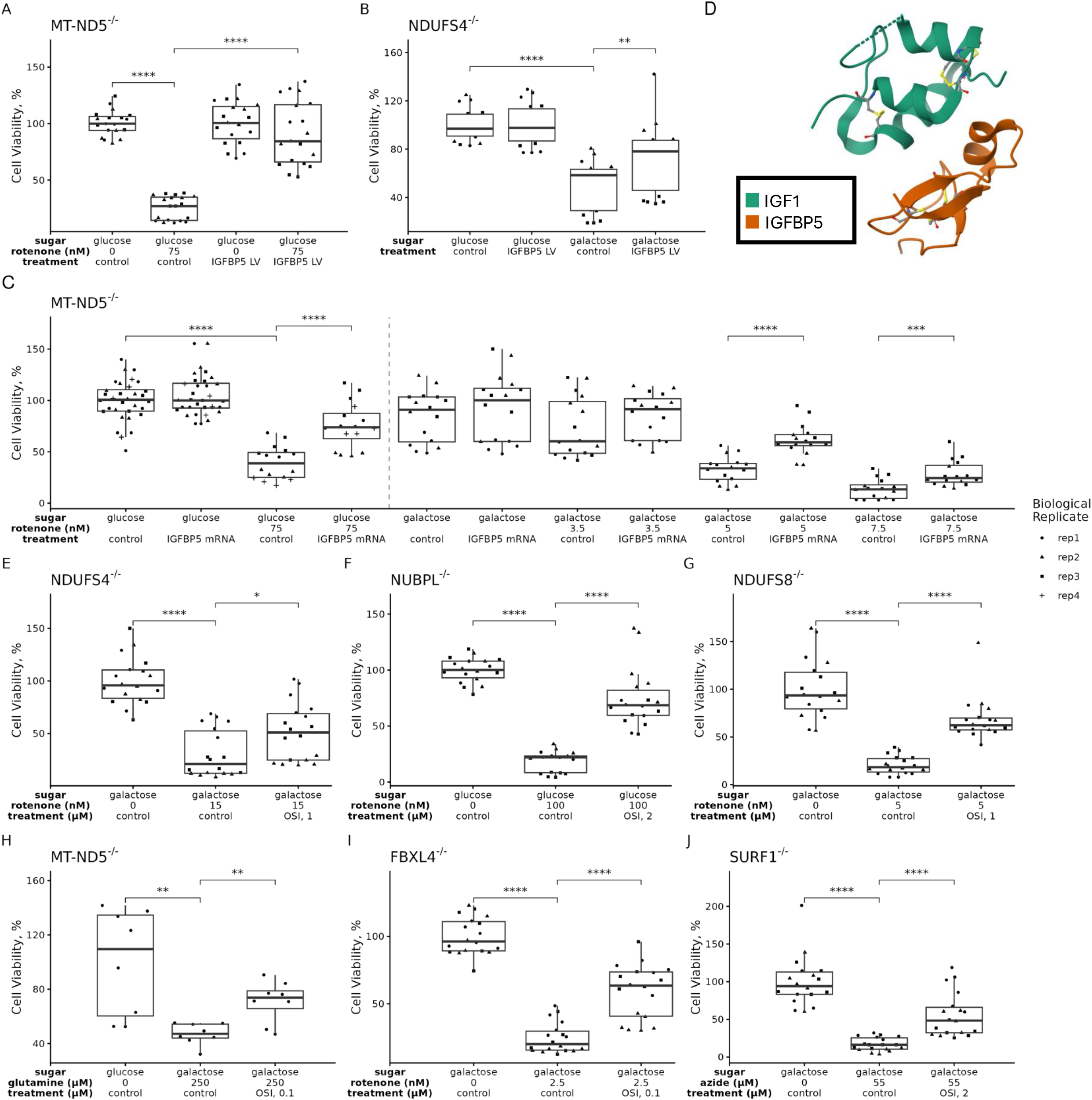
Insulin pathway modulation by IGFPB5 overexpression or IGF1 inhibition rescued mitochondrial disease cell survival under metabolic stress. **(A)** *MT-MD5* fibroblast cell survival was reduced in galactose media with 75 nM rotenone exposure. IGFBP5 overexpression by lentivirus (LV) transfection rescued survival. (B) *NDUFS4^-/-^*fibroblast cells had increased survival after overexpression of IGFBP5 in galactose media. (C) *MT-ND5* fibroblasts had reduced survival in glucose media with high-dose (75 nM) rotenone and in galactose media with low-dose (3.5, 5, or 7.5 nM) rotenone, conditions which were significantly rescued upon IGFBP5 mRNA overexpression at 5 or 7.5 nM rotenone. **(D)** Crystal structure (PDB 1H59) showing direct binding of IGFBP5 (orange) to IGF1 (green) [29]. (E) *NDUFS4^-/-^*fibroblast cells treated with 1 µM of an IGF1 chemical inhibitor, OSI-906, showed significantly increased survival in galactose media with 15 nM rotenone as compared to untreated control cells. (F) *NUBPL^-/-^* cells treated with OSI-906 (2 µM) show increased survival when grown in glucose media with 100 nM rotenone as compared to untreated control cells. (G) *NDUFS8^-/-^* cells treated with OSI-906 (1 µM) showed increase survival, with a stronger effect than treatment with CHX. (H) *MT-ND5* cells in low glutamine media (250 µM) had increased survival in 0.1 µM OSI-906 compared to untreated controls. (I) *FBXL4^-/-^* human PMD fibroblast cells with multiple OXPHOS complex deficiency were evaluated in upon treatment with 1 µM OSI-906, which significantly increased their survival in galactose media with low-dose (2.5 nM) rotenone. **(J)** OSI-906 (2 µM) significantly increased cell survival in *SURF1^-/-^* complex IV deficient patient fibroblasts treated with 55 µM azide. All panels convey results of 72-hour treatment duration, with n=1 biological replicate/condition, and 3 technical replicates, unless otherwise specified.. Cell viability was tested with a t-test with Benjamini-Hochberg for multiple test correction: *, p < 0.05; **, p < 0.01; ***, p < 0.001.

Given our unbiased proteomics finding of significantly altered MAPK pathway expression occurring in mitochondrial disease fibroblasts that was prevented by CHX treatment (**Fig. 2**), together with the known role of IGFBP5 and THPO growth factors in modulating MAPK pathway signaling [32, 33], we performed validation experiments that demonstrated direct support for our prediction that MAPK inhibition would yield therapeutic benefit in PMD. Specifically, we evaluated the effects of a pharmacologic inhibitor of P38 MAPK/ERK activity, talmapimod (TAL), on PMD fibroblast cell viability exposed to a variety of stressor conditions. Indeed, in an *NUBPL^-/-^* complex I deficient patient fibroblast cell line, TAL significantly increased cell survival compared to stressor-only conditions of galactose and 1.8 nM rotenone media (**Fig. 5A**). Similarly, TAL rescued cell viability in an *MT-ND5* fibroblast cell line grown in galactose media with 5 nM rotenone (**Fig. 5B**). In an *NDUFS4^-/-^*complex I disease patient fibroblast cell line, TAL treatment significantly rescued cell viability in 2.5 nM rotenone, fully restoring cell survival back to the levels seen in galactose media alone (**Fig. 5C**). TAL also rescued cell viability in the *NDUFS8^-/-^* complex I disease cell line at both 5 nM rotenone (**Fig. 5D**) and 7.5 nM rotenone (**Fig. 5E**). Combination treatment of talmapimod together with OSI-906 increased the rescue in cell viability over either TAL or OSI-906 alone only in the higher concentration (7.5 nM) rotenone condition (**Fig. 5E**). Collectively, these data demonstrate significant therapeutic benefit on cell survival under metabolic stress in diverse in vitro models of mitochondrial complex I deficiency.

**Fig 5.**
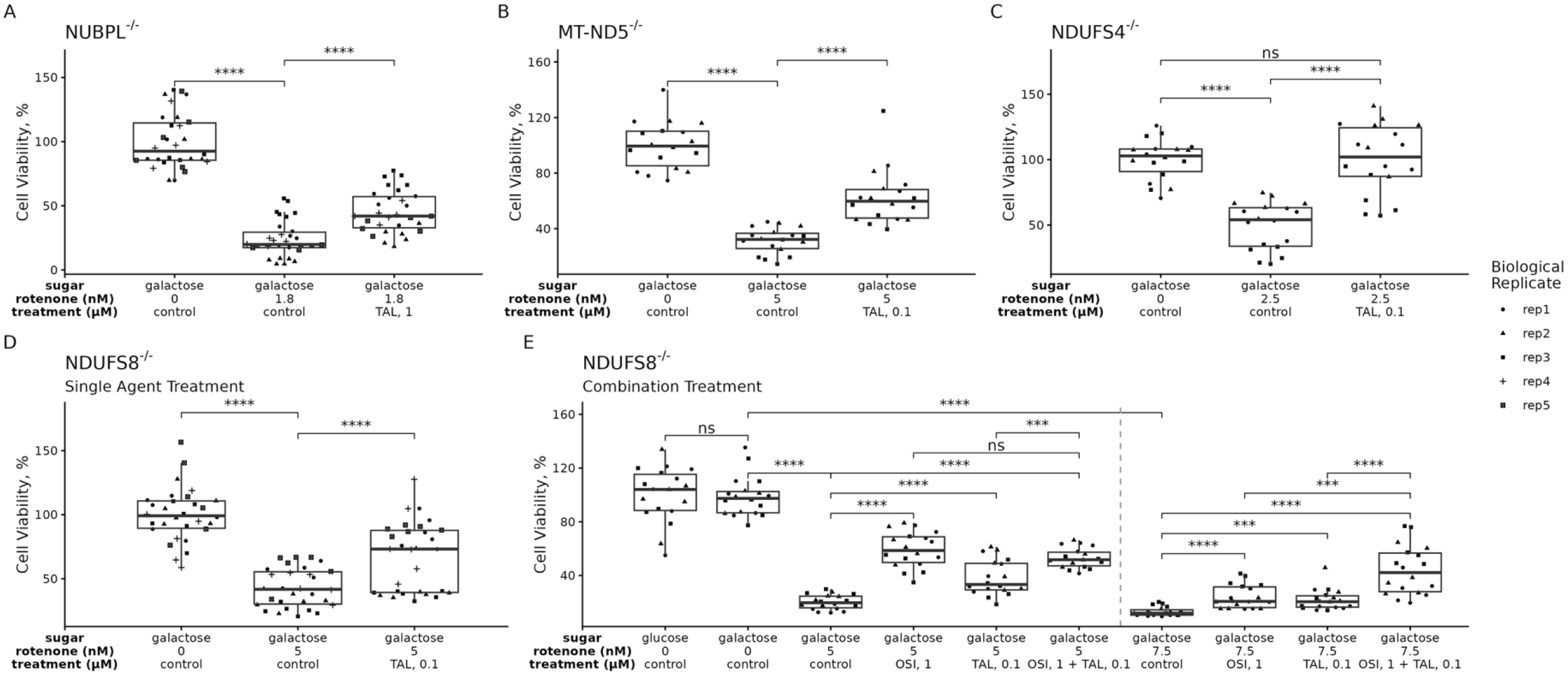
MAPK inhibition rescued mitochondrial disease cell survival. **(A)** *NUBPL^-/-^* cell line had reduced survival after treatment with low-dose (1.8 nM) rotenone in galactose media as compared to culture in galactose media alone. Treatment with a 0.1 µM MAPK inhibitor, talmapimod (TAL), significantly increased cell viability. **(B)** TAL rescued cell viability in a *MT-ND5* cell line after culture in galactose with 5 nM rotenone. **(C)** MAPK inhibition with 0.1 µM TAL in an *NDUFS4^-/-^*cell line rescued cell viability in 2.5 nM rotenone-treated cells back to baseline galactose viability. **(D)** In *NDUFS8^-/-^* cells, TAL alone rescued cell viability in 5 nM rotenone-treated cells. **(E)** Combining TAL and OSI-906 in NDUFS8 cells, both treatments significant rescued cell viability in either 5 or 7.5 nM rotenone. Combining the drugs did not significantly increase rescue under the 5 nM stress condition but did in the higher stress condition at 7.5 nM rotenone. All panels show results of 72 hour treatment duration, with n=1 biological replicates/condition and 3 technical replicates, unless otherwise specified.. Cell viability was tested with a t-test with Benjamini-Hochberg for multiple test correction: *, p < 0.05; **, p < 0.01; ***, p < 0.001.

We next sought to evaluate the whole organismal-level *in vivo* mitochondrial physiology effects of modulating these novel therapeutic PMD targets in an animal model of mitochondrial complex I disease. Specifically, THPO and IGF1 inhibitor effects were evaluated in a well-established *C. elegans gas-1(fc21)* model of complex I mitochondrial disease due to *NDUFS2^-/-^* deficiency [25, 34, 35], using a transgenic worm strain we had generated to contain a knock-in GFP conjugated to hsp-6 (*hsp-6::gfp*) reporter. Hsp-6 is the worm homologue to human Hsp70 that is induced by the mitochondrial unfolded protein response (UPR^mt^) [36], which in turn is activated by mitochondrial stressors that impair OXPHOS capacity, diminish mitochondrial membrane potential, and reduce energy production capacity. In this fluorescent transgenic animal model, the strength of hsp-6::gfp signal induction serves as a direct readout of mitochondrial stress induction [37]. Using this system, the IGF1 and THPO targeted drug therapies identified to have therapeutic benefit in human complex I disease cell lines were studied in *gas-1(fc21)* complex I disease worms imaged 24 hours after treatment initiation at the L4 larval stage. The dynamic range for mitochondrial stress fluorescence induction between wild-type control animals (N2 Bristol worms) and the complex I disease *gas-1(fc21)* model was significant to detect true drug effects (**Fig. 6A**). Further, a previously identified positive control treatment, 2.5 mM *N*-acetylcysteine (NAC) [38], significantly rescued mitochondrial stress induction in *gas-1(fc21)* worms, as measured by the percent reduction in mitochondrial stress (**Fig. 6A-B**). The IGF1 inhibitor OSI-906 at 1 μM (**Fig. 6A**) and the THPO inhibitor ELT at 10μM (**Fig. 6B**) each significantly rescued mitochondrial stress in *gas-1(fc21)* worms, demonstrating U-shaped and linear dose-dependent effects, respectively.

**Fig 6.**
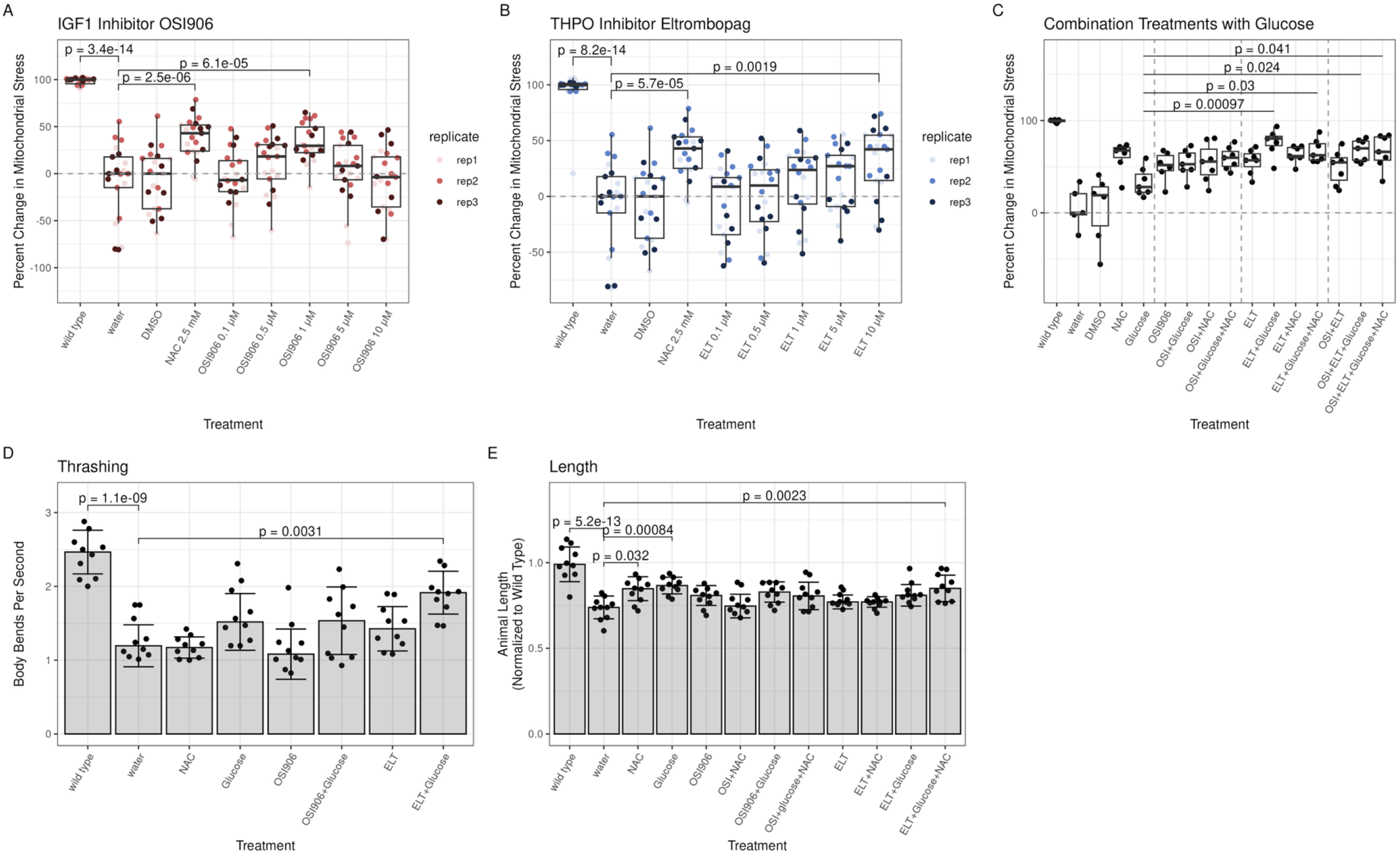
THPO and IGF1 inhibition rescued movement and mitochondrial stress in a *C. elegans NDUFS2^-/-^* model of mitochondrial complex I disease. **(A)** IGF1 inhibitor linsitinib (OSI-906) at 1 μM for 24 hours significantly rescued mitochondrial stress in *gas-1(fc21)* worms, with similar benefit as was seen with assay positive control treatment *N*-acetylcysteine (NAC) at 2.5 mM. By comparison, wild-type (N2 Bristol) worms had 100% less mitochondrial stress than *gas-1(fc21)* worms **(B)** THPO inhibitor Eltrombopag (ELT) at 10 μM for 24 hours significantly rescued mitochondrial stress in *gas-1(fc21) worms*, with similar benefit as was seen with assay positive control treatment NAC at 2.5 mM. **(C)** Combination treatments with the optimal concentrations of 1 μM OSI, 10 μM ELT, and a previously known nutrient treatment in this model of 10 mM glucose. All single- and combination-treatments significantly rescued mitochondrial stress relative to untreated water buffer control (p values not shown). Combinatorial treatments of ELT+glucose, ELT+glucose+NAC, OSI+ELT+glucose, OSI+ELT+Glucose+NAC showed a statistically significant, synergistic, improvement in mitochondrial stress over glucose alone. (D) *C. elegans* thrashing (neuromuscular activity, measured as body bends per second) was significantly rescued with a combination of 1 μM ELT and 10 mM glucose treatments. (E) *C. elegans* body length (growth metric) showed no decrease with either ELT or OSI-906 treatment and was significantly rescue with either known control treatments NAC or glucose as well as by combined ELT and glucose treatments. Across all panels, each circle indicates 1 well, containing a median of between 10 (D-E) or 30 (A-C) animals. All panels show results of 24-hour treatment duration. All differences were tested using ANOVA with a post-hoc Tukey test for all comparisons. In all panels, p values are shown for statistically significant comparisons between treatment groups: *, p < 0.05; **, p < 0.01; ***, p < 0.001.

Combinatorial testing of these individually optimal concentrations of the IGF1 and THPO inhibitors in *gas-1(fc21)* worms showed statistically significant improvement in mitochondrial stress relative to buffer-only *gas1(fc21)* worms (**Fig. 6C** and **Supplemental Table 4** for additional comparison p-values). Importantly, all treatment effects were improved in the setting of glucose treatment [25], as we had previously observed, suggesting glucose itself has therapeutic value in the setting of PMD. However, combinatorial ELT+glucose, ELT+glucose+NAC, OSI+ELT+glucose, OSI+ELT+Glucose+NAC treatments all showed statistically significant rescue over glucose alone (**Fig. 6C**), suggesting nutrient provision together with signaling pathway modulation has synergistic value. Maximal rescue effect of mitochondrial stress induction in *gas-1(fc21)* worms was seen with the combined ELT+glucose treatment.

Gross animal phenotypes were also studied to further assess the *in vivo* therapeutic potential of these novel therapeutic candidates for mitochondrial disease. Neuromuscular activity, as evaluated by thrashing assay, showed significant rescue in *gas-1(fc21)* worms treated with combined 10 μM ELT (THPO inhibitor) and 10 mM glucose (**Fig. 6D**). As THPO and IGF1 are growth factors, worm length was also measured. *Gas-1(fc21)* worms have a linear growth defect relative to wild-type (N2 Bristol) worms (**Fig. 6E**), which was significantly rescued with previously identified antioxidant (NAC) or nutrient (glucose) treatments. Combinatorial treatment with ELT, glucose, and NAC also significantly reduced the linear growth defect of *gas-1(fc21)* worms (**Fig. 6E**).

CHX is a chemotherapeutic agent that we had previously reported to, surprisingly, also have therapeutic benefit in preclinical cell line models of mitochondrial disease [21]. Interestingly, CHX effects in mitochondrial disease cells appeared to be mediated upon *in vitro* proteomics analysis (**Fig. 3**) by THPO and IGF1, which are both growth factors that affect MAPK pathway signaling and have also been associated with increased proliferation in cancer [39, 40]. Therefore, we tested effects of THPO and IGF1 inhibitors on cell viability in two publicly available primary human osteosarcoma cell lines (143B and MG63) relative to hFOB healthy human control osteoblasts and a new cancer cell line 15454-307 that was generated from a lung metastasis from an osteosarcoma patient. Indeed, both THPO inhibition with ELT and as well as IGF1 inhibition with OSI-906 showed dose-dependent killing of all 4 cell lines, with at least one dose showing significantly reduced cell viability in each of the primary and metastatic osteosarcoma cell lines, with the exception of 15454-307 metastatic osteosarcoma cells grown in high glucose that were treated with OSI-906. Whereas these agents’ treatment effects were enhanced in mitochondrial disease cells in the presence of glucose (**Fig. 6**), the reduction of cell viability in osteosarcoma cells was not affected by low (5.6 mM) vs high (17.5 mM) glucose concentration in the culture media (**Fig. 7A**). Combination treatment with both THPO and IGF1 inhibitors at 5 µM ELT and 1 µM OSI-906, respectively, showed greater cytotoxicity relative to untreated cells for hFOB osteoblasts grown in high (17.5 mM) glucose and 15454-307 metastatic osteosarcoma cells grown in low (5.6 mM) glucose. This combination therapy also showed greater cytotoxicity in 143B primary osteosarcoma cells and 15454-307 metastatic osteosarcoma cells grown in low (5.6 mM) glucose, and for hFOB osteoblast control cells grown in high (17.5 mM) glucose, relative to OSI-906 alone (**Fig. 7B**). Whereas the metastatic osteosarcoma line previously showed pan-resistance to a range of therapeutic modalities [41], it is notable that it showed significant response to THPO and ELT inhibition therapies, particularly in combination and low glucose media. Given that osteoblast control cells also showed reduced viability with these interventions, targeting of tumor cells would likely be needed to derive therapeutic benefit while reducing bone cell toxicity from these glucose signaling pathway inhibition therapies in osteosarcoma.

**Fig. 7:**
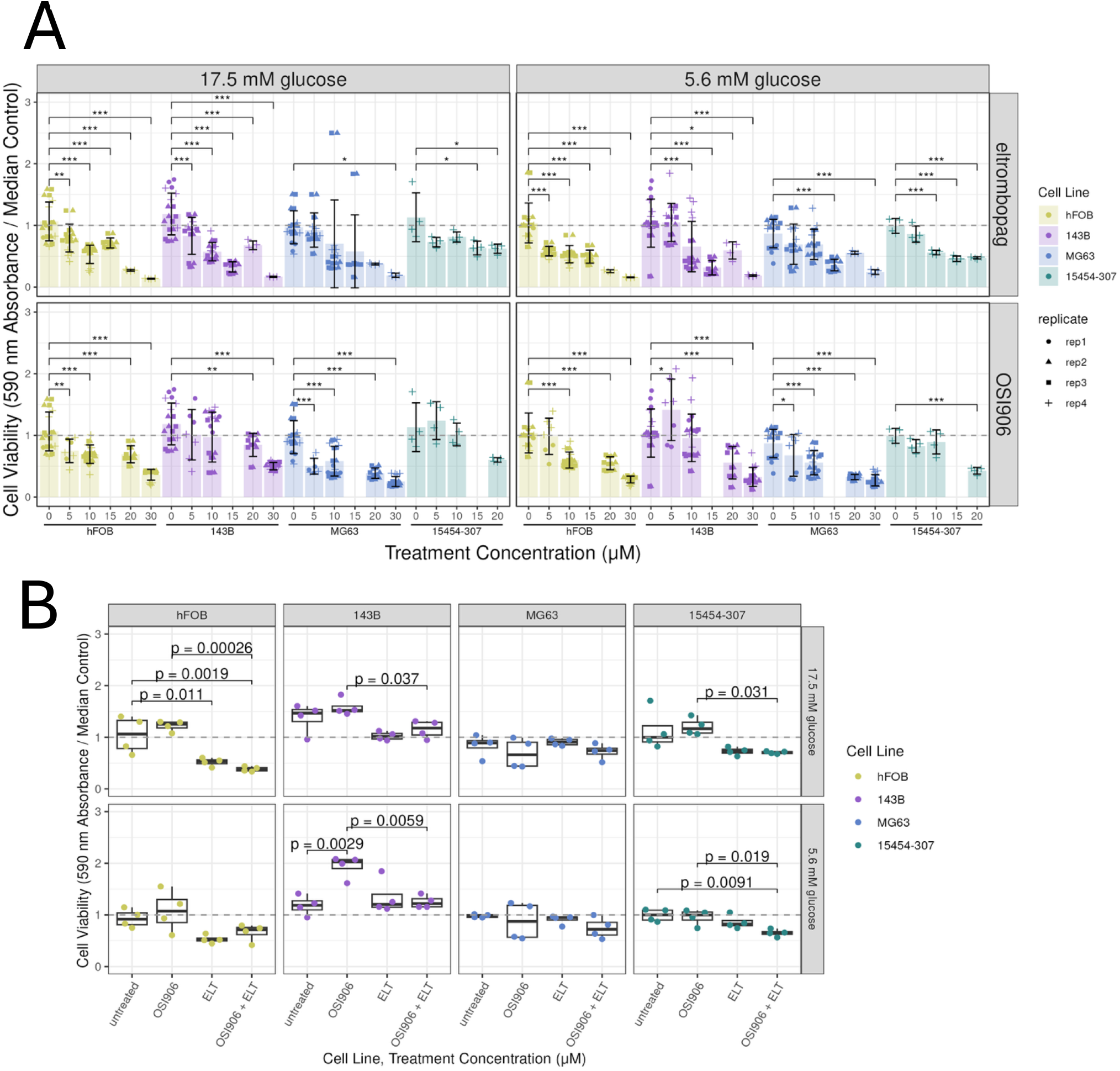
THPO and IGF1 inhibition induced cell death in osteosarcoma cell lines. **(A)** MTT-based cell viability assays were performed to test dose ranges of OSI-906 (linsitinib) and eltrombopag in high glucose (25 mM) and low glucose (5.6 mM) media in a range of cell lines including healthy osteoblast hFOB cells, two primary osteosarcoma cell lines 143B and MG63, and a metastatic osteosarcoma cell line 15454-307. In the heathy osteoblasts and both primary osteosarcoma cell lines, all conditions with both drugs demonstrated a dose-dependent decrease in cell viability and statistically significant killing with at least one dose. Red arrows demarcate doses where selective hypersensitivity in high glucose media was apparent in the primary osteosarcoma lines relative to healthy hFOB osteoblast cells. Metastatic osteosarcoma line 15454-307 showed minimal treatment response in any treatment condition, as measured by minimal reduction in cell viability. **(B)** MTT-based cell viability assays were performed to test effects relative to untreated buffer control of OSI-906 (1 μM), ELT (5 uM), and combinatorial OSI-906 and ELT at equivalent doses. Indeed, combinatorial (1 μM) OSI-906 and (5 μM) ELT significantly decreased cell viability compared to untreated control in healthy hFOB osteoblast cells in high (17.5 mM) glucose media and in 15454-307 metastatic osteosarcoma cells grown in low (5.6 mM) glucose media; no statistically significant difference was seen between combinatorial treatment and treatment with ELT alone. Combinatorial OSI-906 and ELT therapy significantly increased cytotoxicity over OSI-906 alone in 143B primary osteosarcoma cell line, 15454-307 metastatic osteosarcoma cell line, and hFOB osteoblast control cells grown in 17.5 mM glucose.

We sought to further examine the preclinical efficacy of all three drugs (ELT, OSI-906, and TAL) that had been identified by unbiased proteomics analysis in complex I disease cell lines in a wider variety of PMD animal models beyond isolated complex I deficiency. *C. elegans* models previously established in our research laboratory were evaluated with a three-point dose curve (0.1 μM, 1 μM, and 10 μM) of each drug for 24 hours from stage L4 to L4+1 to evaluate their ability to prevent mitochondrial stress (UPR^mt^) induction. In two *C. elegans* RNAi knockdown models of mitochondrial aminoacyl-tRNA synthetases, namely *dars-2* (**Fig. S4A**) and *fars-2* (**Fig. S4B**) [42], none of the three drugs showed significant effect at the level of UPR^mt^ induction. However, the highest dose tested of TAL (10 μM) was toxic in the *dars-2* model, which had previously shown the greatest degree of UPR^mt^ induction of all 19 mitochondrial aminoacyl-tRNA synthetase models studied [42]. By contrast, 100 μM ELT or TAL rescued mitochondrial stress in a novel *slc25a46^-/-^ C. elegans* mutant model (**Fig. S4C**), where *SLC25A46* encodes a mitochondrial outer membrane protein that regulates mitochondrial dynamics. Looking primary human samples, in a fibroblast line from a patient with an *MT-ND1* (m.3985G>A, 66.5% heteroplasmy) variant, we saw a trend towards rescue, although it needs further replication (**Fig. S4D**). We also studied two human Cockayne syndrome patient fibroblast cell lines from affected siblings with identical autosomal recessive pathogenic variants (c.1526+1G>T and c.2800C>A compound heterozygous) in *ERCC6*, a nuclear gene encoding a DNA repair protein in the nucleotide excision repair pathway, but detailed characterization of these patients and cell lines identified complex I deficiency [43]. In fibroblast cells from the more severely affected proband (patient 1), both 0.1 μM OSI and 1 μM TAL increased cell viability over BSO control, while the more mildly affected sibling (patient 2) cell line showed rescue at 1 μM ELT, suggesting these drugs may have differing benefits at different levels of Complex I deficiency (**Fig. S4E**). Collectively, these data suggest that modulation of THPO, IGF1, and MAPK signaling may hold general therapeutic value for complex I deficiency and other complex forms of PMD, but their relative therapeutic benefit at the level of cell survival and mitochondrial stress reduction may vary by specific gene etiology and/or individual patient. A schematic overview depicting the mechanistic cellular effects of these insulin signaling pathway modulation therapies is shown in Fig. 8.

**Figure 8:**
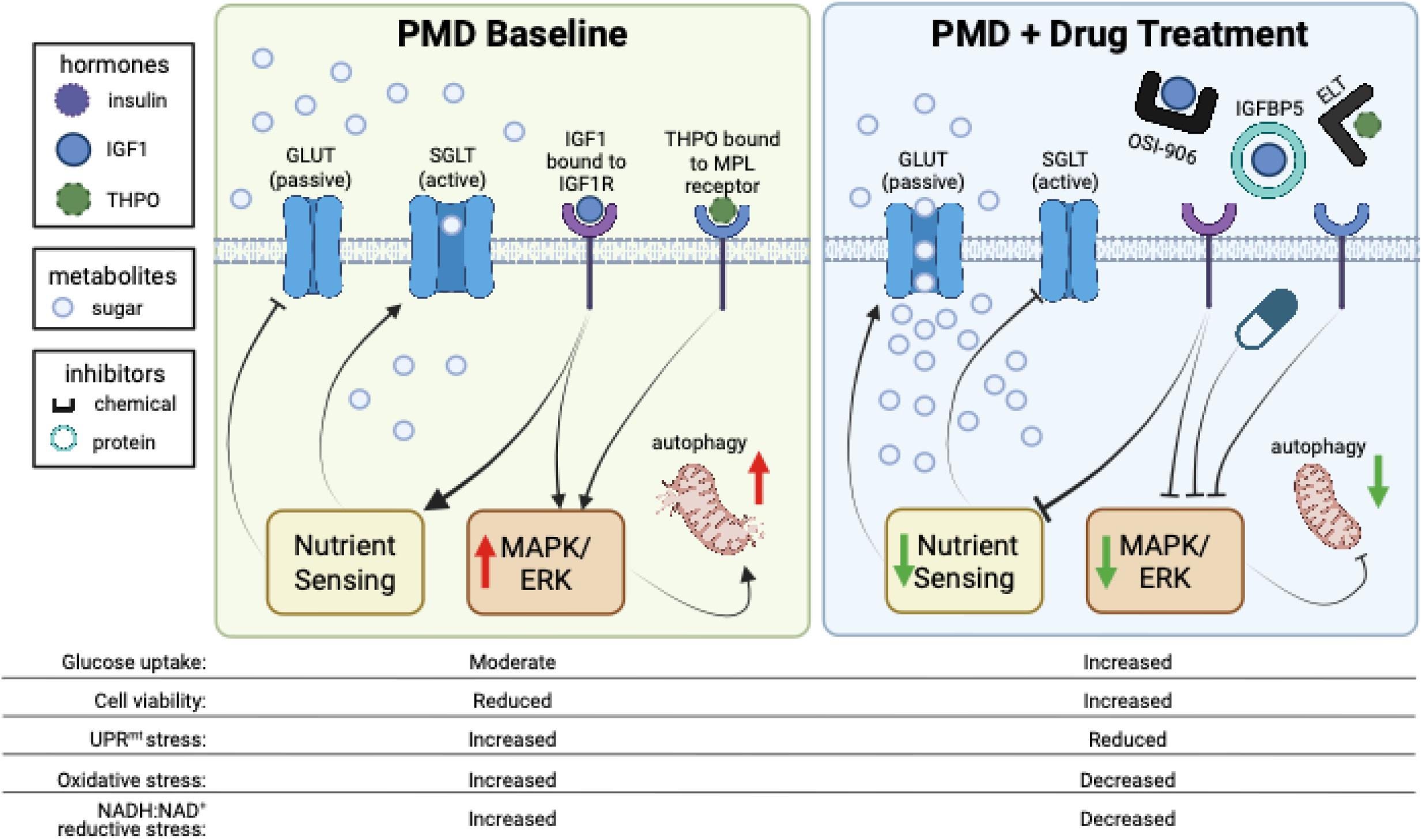
Proposed model underlying the efficacy of IGF1 inhibition in primary mitochondrial disease (PMD). In the absence of any inhibitor, IGF1 binds to its receptor IGF1R, initiating cellular cascades that upregulate both the MAPK and PI3K pathways, wherein the PI3K pathway upregulates cytosolic membrane-bound glucose transporters (enhancing cellular glucose import) and the MAPK pathway upregulates autophagy (cellular recycling). Upon inhibition of IGF1, the PI3K and MAPK pathways are downregulated, limiting glucose import and autophagy. While limited glucose uptake is predicted to be toxic in the setting of mitochondrial disease, our data demonstrate instead that the net effect is to rescue cell viability. Thus, we postulate that glucose uptake is upregulated by IGF1 inhibition through a non-canonical signaling pathway, which together with reduced apoptosis, is therapeutic in PMD and other disorders of mitochondrial dysfunction. Further modulation glucose metabolism, LDHB is involved with regulating intracellular pyruvate-lactate levels and high LDHB downregulates insulin signaling, possibly through NAD^+^-mediated sirtuin activity, providing more evidence we are not invoking the canonical signaling pathways. THPO, similarly to IGF1, upregulates the MAPK pathway upon binding to its receptor, so we hypothesize inhibiting it benefits mitochondrial dysfunction by downregulating autophagy only.

## DISCUSSION

Cycloheximide (CHX) was previously shown in preclinical models to have substantial therapeutic potential for mitochondrial disease [21]. However, as a cytosolic translation inhibitor, CHX causes global changes to cellular regulation with known toxicity in its previous use as an antibiotic or chemotherapeutic agent that [44] precluded its clinical development for PMD [45]. To identify the specific proteins that our prior work suggested to be paradoxically increased upon CHX administration of PMD cells [21], we performed quantitative aptamer-based proteomics to develop a new matrix for SomaLogic quantitation by SOMAscan technology of ∼7,500 proteins using fibroblast cell lines from PMD patients, with subsequent functional validation of differentially expressed targets by CHX treatment performed using a complementary suite of genetic and pharmacologic modulation strategies. Surprisingly, only two differentially expressed gene were identified upon CHX treatment, namely increased expression of IGFBP5 and decreased expression of THPO, together with ERK/MAPK pathway dysregulation,. THPO targeted inhibition both via siRNA knockdown and small molecule chemical inhibition (eltrombopag), significantly rescued survival under metabolic stress in fibroblast cell lines from multiple PMD patients. IGFBP5 overexpression by lentiviral or mRNA approaches likewise rescued PMD cell survival, as did pharmacologic inhibition (OSI-906) of its known direct binding partner, IGF1, which modulates the insulin signaling system. THPO and IGF1 inhibition were functionally validated to significantly rescue mitochondrial stress induction, linear growth deficiency, and neuromuscular thrashing activity impairment in the well-established *C. elegans gas-1(fc21) ndufs2^-/-^* animal model of mitochondrial complex I disease. Direct p38 MAPK inhibition itself using a small molecule, talmapimod, also significantly improved fibroblast cell survival in diverse PMD disorders. These therapies were shown to have enhanced efficacy when given in combination with high glucose, which we have previously demonstrated to have significant therapeutic value in diverse models of PMD [24, 25]. Additionally, modulation of MAPK pathway signaling, which is known to be dysregulated in cancer, by THPO and IGF1 inhibition synergistically acted to significantly kill primary and metastatic osteosarcoma cells, although with a narrow therapeutic window over control osteoblast cells that suggests a potential need for tumor-specific targeting.

When considering the presumed mechanism of enhancing insulin signaling to increase cellular uptake of glucose and the therapeutic role of glucose in PMD, IGF1 inhibition as a therapeutic strategy in mitochondrial disease appears counterintuitive. Indeed, the canonical effect of IGF1 is to bind to its receptor, IGF1R, which activates signaling cascades that ultimately upregulate the plasma membrane glucose transporter, GLUT1-10 (aka SLC2A1-10) (**Fig. 8**) [46]. According to this classical model, IGF1 inhibition should downregulate glucose import into cells by downregulating their transporters, which should be highly toxic for nutrient-dependent mitochondrial disease cell lines in which we have shown that glucose itself has significant therapeutic value [24, 25]. Nonetheless, the results of the proteomics and subsequent functional validation studies in cell lines and *C. elegans* models detailed here demonstrate the clear therapeutic value of IGF1 inhibition in PMD. Further, counter to classical insulin signaling pathway dogma, Meriin et al recently demonstrated that inhibitors of RNA translation, like CHX, actually stimulate glucose receptor translocation to the plasma membrane and increase glucose uptake, independent of the canonical pathways [47]. Therefore, we postulate that CHX treatment, IGBP5 overexpression, or IGF1 inhibition similarly increase cellular uptake of glucose (and other carbon sources including galactose that utilize GLUT transport [48]) likely through the same yet-to-be discovered non-canonical pathway, allowing PMD cells that have impaired OXPHOS capacity to utilize glycolysis to more efficiently generate biochemical energy in the chemical form of ATP. Indeed, *C. elegans* animal model studies clearly demonstrated the increased efficacy of both THPO and IGF1 inhibition when given in combination with glucose. These results provide evidence suggestive of this hypothesis that the net effect of inhibiting THPO and IGF1 signaling is to increase glucose uptake and utilization to yield cellular energy.

Autophagy inhibition is likely a second mechanistic benefit by THPO or IGF1 inhibition in the context of mitochondrial disease via decreasing MAPK signaling [27, 32]. Our previous *in vitro* studies previously showed autophagy to be significantly upregulated in the setting of mitochondrial respiratory chain deficiency, which was prevented by CHX [21]. Here, proteomic analysis of CHX effects in complex I disease fibroblast cells demonstrated it to prevent induction of the MAPK pathway. As IGF1 and THPO are both known activators of the MAPK pathway [27, 32], inhibiting them and therefore MAPK provides dual-pronged cellular benefits for PMD, both increasing the supply of glucose and preventing dysfunctional induction of autophagy.

Direct MAPK pharmacologic inhibition with talmapimod supports [49, 50] this therapeutic model, as improved cell survival was seen under metabolic stressor conditions in three mitochondrial complex I disease patient fibroblast cell line models. Interestingly, MAPK inhibition was also previously reported to have therapeutic value in a distinct form of PMD, namely Friedrich Ataxia [51]. Together, these insights suggest future clinical research to advance development of CNS-penetrant MAPK inhibitors may be a valuable path to consider in PMD.

Cancer is widely viewed as having PMD-like disease properties, as they display the well-known Warburg-effect, a reliance on glycolysis even in the presence of sufficient oxygen [52], similar to how PMD cells rely on glycolysis due to their defective mitochondria. However, in recent years, more work has come to endorse a more nuanced approach, both glycolysis and oxidative phosphorylation [53] and recent work in our lab has shown that osteosarcoma is strongly dependent on its mitochondrial function for survival [41]. As our mechanistic hypothesis is that these targets are improving function in PMD by directing energy metabolism away from the mitochondria towards glycolysis, we asked if doing the same in mitochondrial-reliant cancer would have a killing effect, which, remarkably, they did. These findings are similar to the dual-indication effects of CHX, which we recently reported to be cytotoxic in cancer cells while improving cell viability in a glucose-dependent fashion in mitochondrial disease cells [41]. Interestingly, these glucose signaling pathway therapies appeared to show greater cytotoxicity in control osteoblasts than in control fibroblasts, speaking to the need to further evaluate tissue-specific adverse effects of doses advanced toward clinical trials. Further, it is notable that a highly treatment recalcitrant pulmonary metastasis derived osteosarcoma line showed significant cell death in response to THPO and ELT inhibition therapies, particularly in combination and low glucose media, as this line previously showed pan-resistance to a range of therapeutic modalities [41] and metastatic osteosarcoma has significant mortality. Given that modulation of these protein targets may be amenable to a more targeted tumor treatment approach, targeting these therapies in cancers, such as osteosarcoma that was studied here, are likely to yield the benefit of tumor cytotoxicity without the broader adverse sequelae that occur in cancer patients treated with CHX due to its many off-target effects.

Overall, IGFBP5/IGF1, THPO, and MAPK are demonstrated by this work to represent (1) novel pathogenic mediators of mitochondrial respiratory chain disease, (2) the precise protein and pathway (MAPK/ERK, insulin-stimulated glucose signaling) mediators mechanistically underlying the therapeutic benefit of CHX in preclinical mitochondrial disease models, and (3) novel therapeutic targets whose specific modulation have demonstrated, and synergistic, survival and functional benefits evident across evolutionarily distinct human cell and *C. elegans* preclinical models of diverse mitochondrial diseases. Furthermore, nutrients in the form of glucose, alone or together with antioxidant therapy in the form of NAC, showed further synergy with THPO and IGF1 signaling pharmacologic inhibition in PMD. Future clinical trials are warranted to advance genetic and/or pharmacologic strategies that selectively activate IGFBP5 (such as by mRNA or gene therapies) or inhibit IGF1, THPO, or MAPK, with broad therapeutic potential across a wide range of individually rare genetic causes of PMD, as well as to consider these as adjuvant therapies for metastatic cancers such as osteosarcoma.

## MATERIALS AND METHODS

### Sex as a biological variable

Sex was not considered as a variable in this paper. However, mitochondrial patient fibroblasts were obtained from 3 male (*NDUFS1^-/-^*, *MT-ND1*, *ERCC6^-/-^* patient 1) and 7 female patients (*NDUFS8^-/-^*, *NDUFS4^-/-^*, *MT-ND5*, *NUBPL^-/-^*, *FBXL4^-/-^*, *SURF1^-/-^*, *ERCC6^-/-^*patient 2). For the proteomics, matched samples from their mothers were used to ensure a matching mitochondrial background in the absence of disease. As *C. elegans* are hermaphrodites, sex-specific effects are not applicable. The findings are expected to and have been equally effective in preclinical models of both sexes.

### Study approval

All mitochondrial patient samples were collected at Children’s Hospital of Philadelphia under IRB #08-6177 (PI: MJF).

#### Sample collection for SomaLogic DNA aptamer-based proteomics

Mitochondrial disease patient samples and healthy controls were collected and generated at the Children’s Hospital of Philadelphia (IRB #08-6177). Cells were grown in 5.6 mM glucose media in a 6-well plate until they reached 80% confluence. Sample was then collected for glucose control, and remaining wells treated with 100 nM rotenone, 3.6 μM cycloheximide, or a combination of both. Cells were then harvested at either 4 or 24 hours after treatment. To extract protein, cells were washed 3 times with DPBs and then treated with 300 μl lysis buffer (RIPA buffer, protease inhibitor, 25 mM Tris, 150 mM NaCl, 1% NP-40, 1% sodium deoxycholate, 0,1% SDS). Then cell lysates were scraped into microfuge tubes and centrifuged for 5 min at 14,000 x g. Total protein was measured using the Pierce BCA method and then aliquoted in 75μl (15 μg) aliquots at a concentration of 200 μg/ml in accordance with SomaLogic’s sample requirements. Samples were frozen at −80°C before being shipped to SomaLogic for protein quantification.

#### Protein Quantification

SomaLogic executed the SOMAscan assay, which has been extensively described previously [54, 55]. Briefly, high-affinity DNA aptamers were developed from random sequence libraries through the Systematic Evolution of Ligands by Exponential enrichment (SELEX) progress, which has successive rounds of *in vitro* selection and enrichment to identify the most selective sequences with slow off-rates. These aptamers are functionalized on the 5’ end a fluorescent tag, a photocleavable linker, and biotin, the last of which allows the aptamers to be attach ed streptavidin beads. The protein sample and beads are mixed, allowing proteins to bind to the aptamers and then washed. After washing, samples are exposed to ultraviolet light to break the photocleavable linker, releasing the aptamer-protein complexes, which are then recaptured on a second set of beads for further purification. After washing again, the aptamers are eluted under denaturing conditions and quantified via DNA microarray, quantifying the protein values as the relative fluorescence signals (RFUs) of their corresponding aptamers [55].

Fluorescence values were normalized by SomaLogic following their standard pipeline to remove systematic biases in the assay on a per-plate basis. Briefly, blank hybridization controls without protein are included to account for background and to calculate a scaling factor that is applied to all other spots on the microarray. Calibration controls, specific to the matrix being used (Ex. plasma, serum, fibroblasts), are invariant and the median is taken to calculate a calibration scaling factor for all aptamers and plates. Plate scaling is also performed to normalize signal across plates. Some aptamers are also repeated with dilution and a scale factor for each dilution group is calculated. With these scaling factors, adaptive normalization through maximum likelihood, is used to normalization the data. SomaLogic has previously demonstrated the efficacy of the methods to increase reproducibility and reduce systematic biases [55]

#### Protein Analysis

Differential proteomics was analyzed using custom R scripts [56, 57] following SomaLogic’s best practices. Differences in protein expression between groups were tested with ANOVA followed by a post-hoc Tukey test. Pathway analysis was done using WebGestaltR, using their network topology-based analysis method with BioGRID’s protein-protein interaction network as the reference data [58]. Visualizations were made using ggplot2 [59] with the addition of treemapify for pathway analysis results [60] and the figure was assembled using patchwork [61].

#### Mitochondrial Disease Fibroblast maintenance

Human mitochondrial disease or health control fibroblasts were maintained in 10% FBS-5.6mM glucose DMEM-15uM uridine at 37C 5% CO2.

#### Mitochondrial Disease Fibroblast MTT Assay

Mitochondrial disease patient samples and healthy controls were collected and generated at the Children’s Hospital of Philadelphia (IRB #08-6177). Cells were seeded with 100ul of 5k cells/well into a 96-well plate (Corning #3585) with DMEM (5.6mM glucose)-10% FBS-15uM urinine-2mM glutamine and cells were cultured in 37C, 5% CO2 incubator for overnight. After removing medium, 100ul (for 72h treatment) or 150ul (for 6 days treatment) fresh selected medium with or without drugs were added to each well. Cells were kept in incubator for 72h or 6 days. After removing medium, 100ul/well MTT solution (0.45mg/ml MTT (3-(4,5-Dimethylthiazol-2-yl)-2,5-Diphenyltetrazolium Bromide, Thermo Fisher # M6494) was prepared with DMEM by 5mg/ml MTT Stock solution (in PBS)) was added to each well, incubating for 4 hours at 37C, then 140ul/well DMSO was added to each well. After waiting for 15 minutes at room temperature, the absorbance was measured with a 590nm excitation in microplate reader

### siRNA Knockdown Assay

10k cells/well were seeded into 96 well plate with growth medium. Next day Opti-MEM medium (5ul/well), Lipofectamine 3000 Reagent (0.3ul/well) and 10nM siRNA (THPO siRNA, Thermo Fisher #s14114) were prepared. Mixing Opti-MEM and Lipofectamine, waiting for 5 minutes, then mixing mixture of Opit-MEM-Lipofectamine and siRNA solution, waiting for 20 minutes at room temperature. Add 10ul/well mixtures into 96 well cells in 90ul/well fresh growing media. Incubate cells for 48h, then test cell viability with MTT assay.

#### IGFBP5 overexpression by lentivirus-igfbp5 transfection

75k cells/well were seeded into 6 well plate with growth medium DMEM (5.6mM glucose-10% FBS-15uM uridine. IGFBP5-lentivirus (ABM #LV7-24325061) was prepared to 10^5^TU/ml – 10ug/ml polybrene with growth medium. Medium was removed from 6 well plate with the cultured cells, and then 1.5ml/well of igfbp5-lentivirus-polybrene solution was add to each well. Cells were cultured for 48h, then split into a T25 flask. Two days later, 1ug/ml purimycin was added for selection.

#### IGFBP5 mRNA overexpression

Patient fibroblasts (passage number below 6) were grown in 5.6mM glucose DMEM-10% FBS-15uM uridine medium in T150 flasks at 37 °C in a 5% CO2 incubator to 90% confluency, then 5-10k cells/well were split into a flat bottom 96 well plate with the same growing medium and incubated for an additional 24 hours. Media was then removed, and replaced with 100 μl growing medium with 0, 10, 25, 50, 100, or 200 ng/well of IGFBP5-LNP-mRNA (University of Pennsylvania, see **Fig S5** for sequence). After 24 hours of incubation, the media was removed, and 100 μl DMEM-10% FBS with 5.6mM glucose or 2.8mM glucose-5mM galactose or 10mM galactose with or without different doses of rotenone was added. After 72 hours of incubation, cell viability was read in 96 well plate with MTT following the protocol as above.

#### *C. elegans* strains and maintenance

C. elegans wild-type N2 Bristol worms, SJ4100 (*zcIs13[hsp-6p::gfp+ lin-15(+)]*), VS21 (*hjSi20 [myo-2p::mCherry::unc-54 3’UTR]*) and CW152 (*gas-1(fc21)*) were obtained from the Caenorhabditis Genetics Center (CGC). Two *C. elegans* fluorescent knock-in strains were made, MJF2/N2 (*hsp-6p::gfp + myo-2p::mCherry*) and MJF3 (*gas-1(fc21)* (*hsp-6p::gfp + myo-2p::mCherry*)). Animals were maintained at 20 °C on nematode growth media (NGM) OP50 plates (20X concentration of OP50 *E. coli* on 100 mm diameter plates) unless stated otherwise.

#### Worm synchronization

Synchronous populations of *C. elegans* were obtained by bleaching. Worms were collected from the 100 mm NGM plate with 4 mL S. Basal (23.4 g NaCl, 4 g K_2_HPO4, 25 g KH_2_PO4, 4 ml cholesterol (5 mg/ml in ethanol), 4 L milliQ water, pH = 7). 1 mL 100% bleach and 500µL NaOH 5M was added to the worm suspension and the tubes were mixed for 2-3 min until no visible worm body was present. The solution was centrifuged at 182 rcf for 1 min, the supernatant was immediately discarded, and the pellet was washed twice with 10 mL S. Basal. The embryos were then seeded on 100 mm NGM plates seeded with OP50 bacteria and incubated at 20°C.

#### *C. elegans* Mitochondrial Stress Assay

Worm embryos were obtained by bleaching and plated on 100 mm NGM plates seeded with OP50 bacteria for 3 days. Adult stage L4 worms were then collected with 5 mL S. Basal supplemented with OP50 (OD600 of 0.7). The worm suspension was diluted to a concentration of 1 worm/µL, and 50µL per well was dispensed into a 384 well plate (ThermoFisherScientific, 142761). Drug treatment was added to each well and the plate was incubated at 20°C for 24h.

Mitochondrial stress was measured using a CellInsight CX5 (Thermo Scientific) using a protocol adapted from previous high-throughput *C. elegans* screening (PMID: 21103396). Briefly, SpotDetector was used to focus the red channel and a 20% exposure was sufficient to detect all the worms in the wells, an exposure setting of 40% was used for the green channel and 10% was used for the brightfield image, these exposure settings were set for untreated MJF3 well. The red channel was used to count the number of worms in each well, the green channel was set to a threshold of 1800 pixel intensity and, using spot total intensity, was used to measure the total green fluorescence of each well.

#### C. elegans Thrashing

Synchronized L4 worms were transferred to 3.5 cm NGM plates containing the desired concentration of drugs, positive control or buffer control, and incubated at 20°C for 24h. 1 mL room temperature S. Basal was then added and the worm swim activity was recorded for 10s using a Basler USB Camera (model #108014) with a KOWA industrial lens 75mm/F2.5. The FIJI plugin wrMTrck [62] was used to measure the body bends per second of each individual *C. elegans* and for three biological replicates [63].

#### *C. elegans* Mitochondrial Stress Data Normalization and Analysis

Data analysis was conducted using custom R scripts [56, 57]. For the mitochondrial stress assay, for each well, the GFP was normalized as the percent reduction in stress relative to the plate controls. Background, healthy animal fluorescence, was subtracted from all values and then percent change between the median disease fluorescence and the well was calculated for each well.

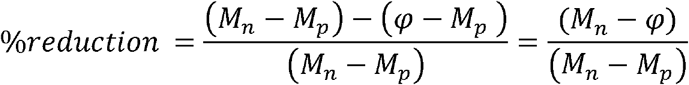

Where n is the negative control *gas-1*, p I s the positive control N2, *n* is the median of the control for the given the subscript for the plate and is the green fluorescence intensity from the well.

For both the mitochondrial stress assay and the thrashing assay, differences between conditions were tested with an ANOVA followed by a post-hoc Tukey test and visualized using ggplot2 [59] and assembled into a single figure using patchwork [61].

#### Osteosarcoma Cell Lines and Reagents

Human fetal osteoblast hFOB 1.19 (CVCL_3708, hFOB), 143B, MG-63 cell lines were obtained from the American Type Culture Collection (ATCC). The 15454 cell line was generated from a pulmonary metastasis from an osteosarcoma patient isolated at Children’s Hospital of Philadelphia (IRB #18-015454). All cell lines were maintained in a CO_2_ incubator at 37°C, 5% CO_2_. All cell lines were maintained in DMEM:F12 (Gibco, 11330) supplemented with 10% FBS. Cell lines were routinely monitored until 70%-80% confluency was observed then were appropriately seeded for the experiments. All cell lines reported a negative mycoplasma result. Eltrombopag Olamine and Linsitinib were obtained from Sellekchem, (S2229 and S1091, respectively).

#### Osteosarcoma MTT Assay

All cell lines were plated in a 96-well plate format (2,500 cells/100 µL/well) in DMEM:F12 media then placed in a CO_2_ incubator (37°C, 5% CO_2_). The next day, each well was aspirated then washed twice with dPBS. Then, one of the following medias was used, either DMEM:F12 media (supplemented with 10% FBS) or DMEM-low glucose (Gibco, 11885) supplemented with 10% FBS. The appropriate concentrations of Eltrombopag Olamine or Linsitinib were added to each well then placed into CO_2_ incubator for 72 hours. After 72 hours, added 50 µL MTT reagent (Abcam, Ab211091) and 50 µL serum free media to each well then placed back into the CO_2_ incubator for 3 hours. After 3 hours, added 150 µL MTT solvent (Abcam, Ab211091) to each well. Next, each plate was wrapped in aluminum foil and placed on an orbital shaker for 15 minutes. Next, each plate was read on a multi-well plate reader at OD = 590 nM, along with a reference wavelength, OD = 670 nM.

#### Osteosarcoma Statistical Analyses

Data was analyzed using custom R scripts [56, 57]. Normalization was performed by taking the ratio of the MTT fluorescence value of each well to the median of the untreated control for each cell line. Differences between conditions were test with an ANOVA followed by a post-hoc Tukey test and visualized using ggplot2 [59] with colors from the simplecolors R package [64] and individual plots assembled using patchwork [61].

## Data availability

All data is deposited in PRIDE

## Supporting information

Supplemental Table 1 Differential proteomics abundance

Supplemental Table 2 Proteomics pathway analysis

## ACKNOWLEDGMENTS

The authors are grateful to SomaLogic Proteomics for their generous in-kind research contributions to this study including performance of proteomics data generation on fibroblast cell lines; Rarefy Therapeutics for in-kind provision of Talmapimod under collaborative research agreement; and to Adam Resnick, PhD, and Jessica Foster, MD, for helpful conversations and guidance on designing mRNA overexpression therapy.

## AUTHOR CONTRIBUTIONS

Conceptualization: MJF, NJ, SD (Dugar); Data curation: KK; Data analysis: KK, MP, MJF; Funding acquisition: MJF; Investigation: MP, CR, NW, VM, NW, MK, SD (Dharaskar), SH, APS, DI; Proteomics data generation oversight: NJ; Visualization: KK; Writing – original draft: KK, MP; Writing – review and editing: KK, MP, VA, MJF

