## Supplemental figures for "Targeted modulation of IGFBP5/IGF1, THPO, and P38 MAPK signaling are potent therapeutic strategies generalizable for mitochondrial respiratory chain disease and osteosarcoma"

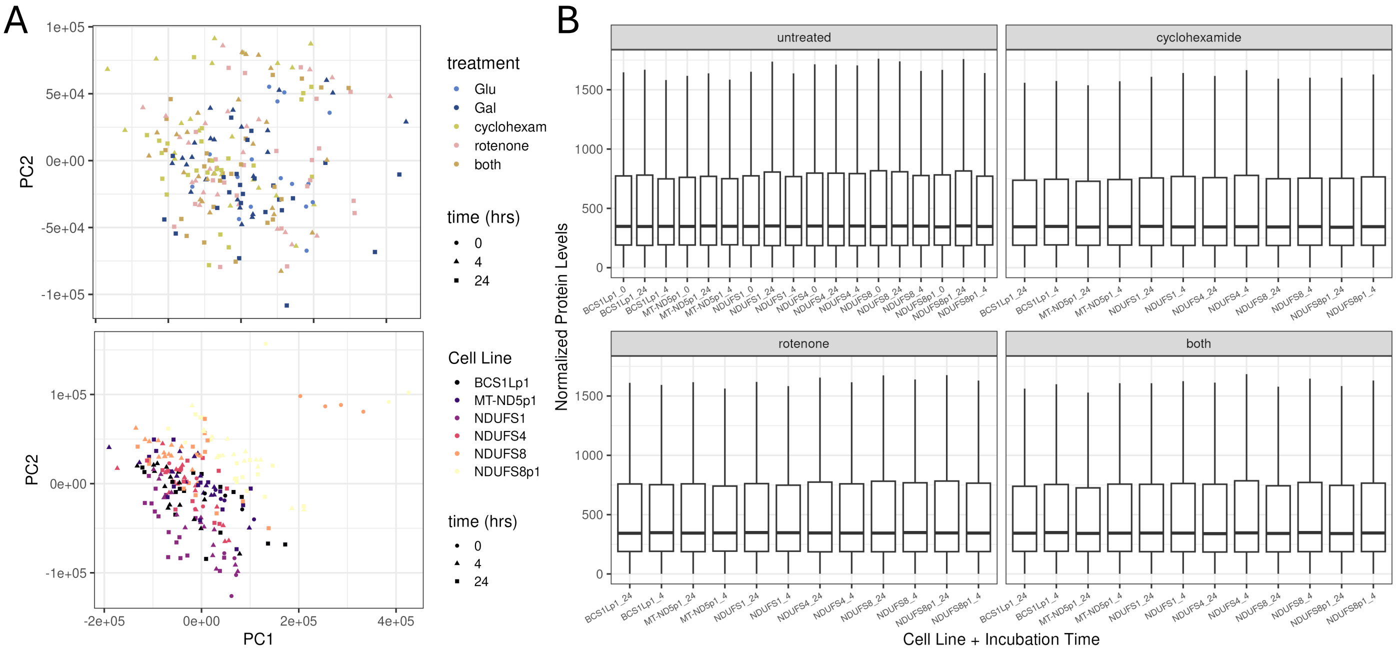


**Figure S1. Quality Control Proteomics Results.** (**A**) Principal Component Analysis (PCA) of all samples. The same PCA is shown twice, once colored by the sample treatment and once colored by the cell line identity. The NDUFS8 (autosomal recessive disease patient) and NDUFS8p1 (heterozygous carrier parent) cell lines were both excluded in this analysis due to technical concerns of their respective gene variant status, and they showed a clustering effect in the PCA. Generally, PCA results did not cluster by treatment but by cell line identity. (**B**) Normalized protein distribution values were equivalent across all samples, as expected.  Sample IDs are shown on the X axis, where affected PMD patient cell lines are indicated by gene name and healthy parents are indicated with the annotation “P1”.

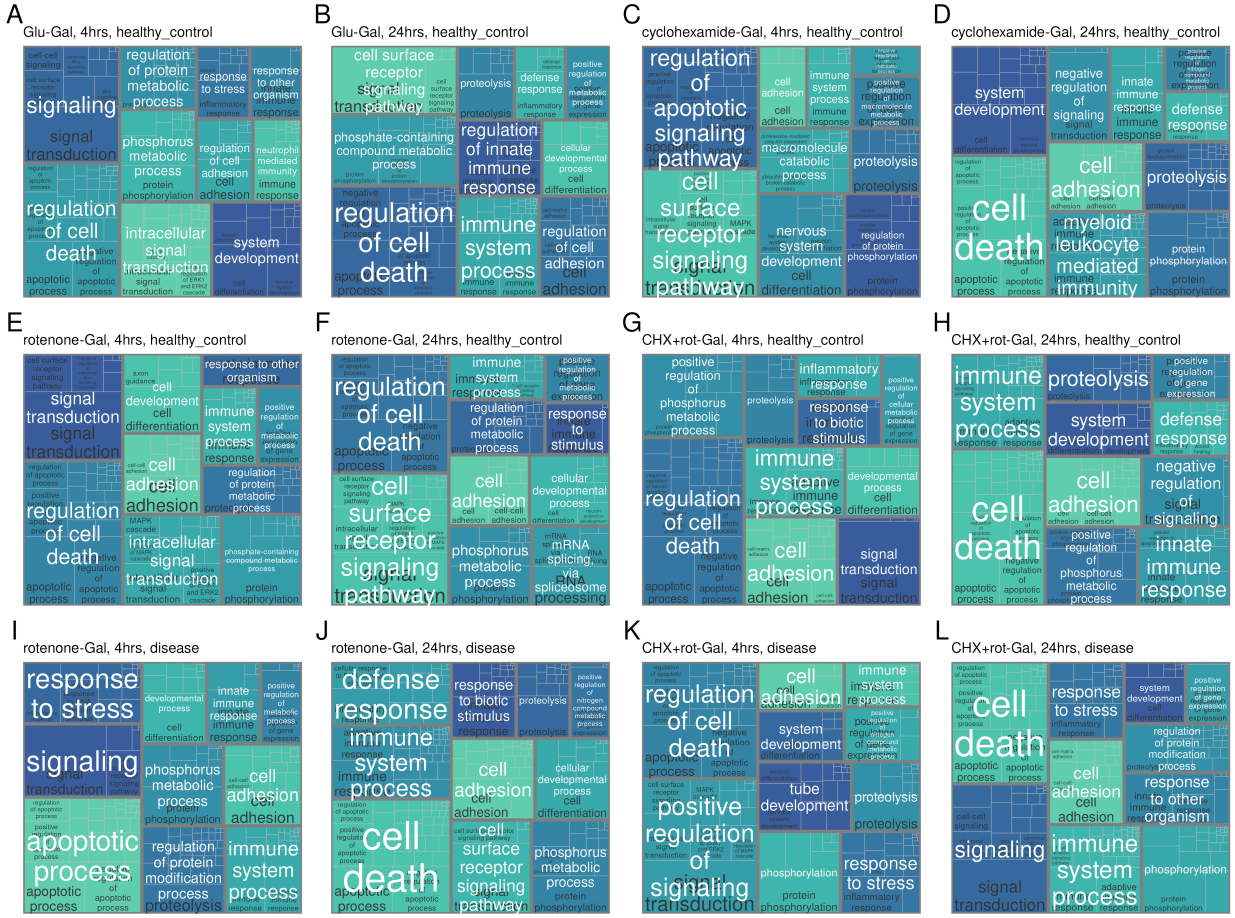


**Figure S2. Pathway analysis of proteomic profiling in all other tested media and treatment combinations in complex I disease and healthy control cell lines.** Comparative **a**nalyses of (**A**) Healthy cell lines, glucose vs galactose at 4 hours, (**B**) Healthy cell lines, glucose vs galactose at 24 hours, (**C**) Healthy cell lines, cycloheximide vs galactose at 4 hours, (**D**) Healthy cell lines, cycloheximide vs galactose at 24 hours, (**E**) Healthy cell lines, rotenone vs galactose at 4 hours, (**F**) Healthy cell lines, rotenone vs galactose at 24 hours, (**G**) Healthy cell lines, combination treatment cycloheximide and galactose at 4 hours, (**H**) Healthy cell lines, combination treatment cycloheximide and galactose at 24 hours, (**I**) Complex I disease cell lines, rotenone at 4 hours, (**J**) Complex I disease cell lines, rotenone at 24 hours, (**K**) Complex I disease cell lines, combination treatment cycloheximide and rotenone at 4 hours, (**L**) Complex I disease cell lines, combination treatment cycloheximide and rotenone at 24 hours.

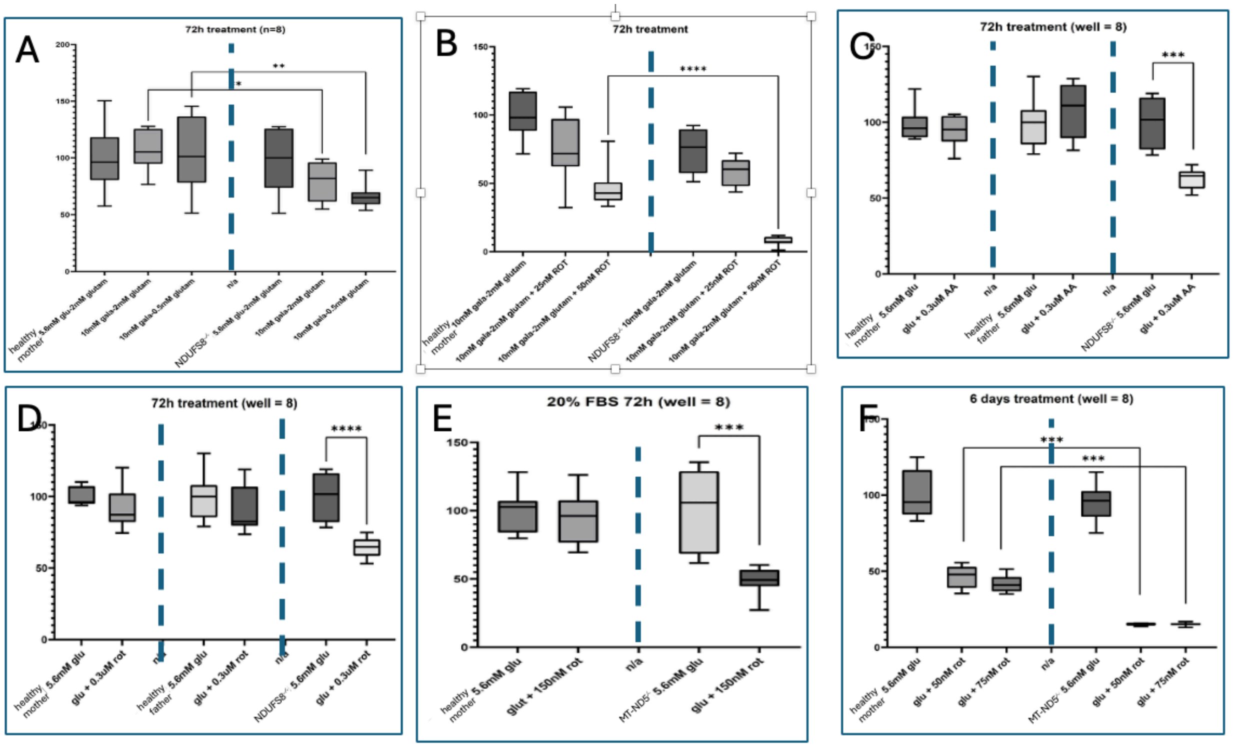


**Figure S3. Experimental analyses of culture conditions for cell viability assays** (A-D) parents of NDUFS8$^{-/-}$ patient and NDUFS8$^{-/-}$ patient (E-F) mother of MT-ND5$^{-/-}$ patient and MT-ND5$^{-/-}$ patient

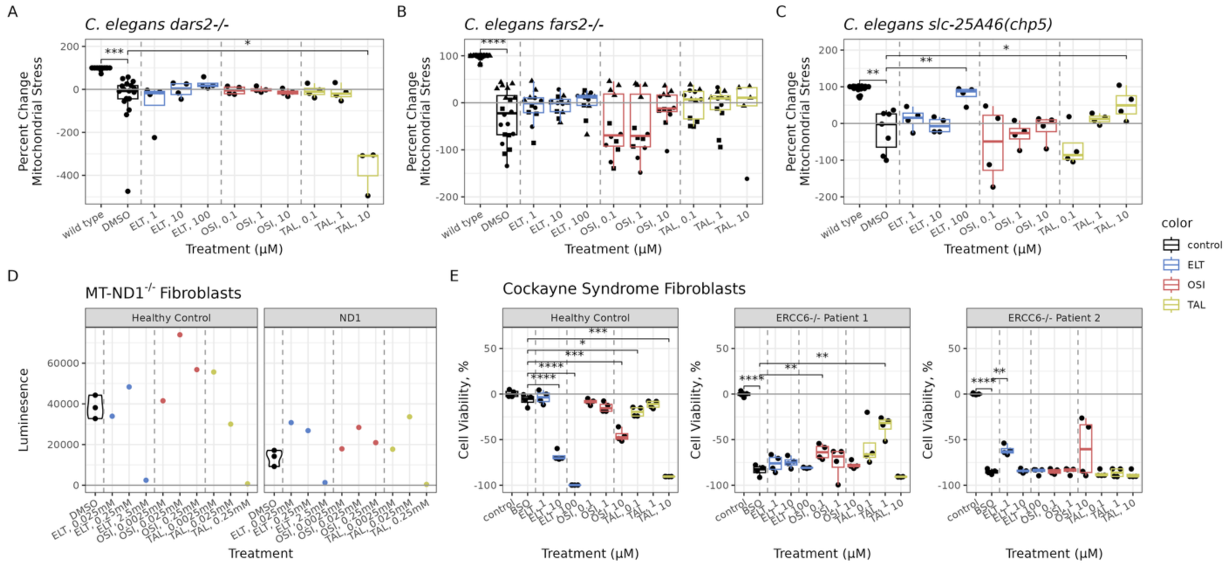


**Figure S4. THPO, IGF1, and MAPK dose curves in other models of mitochondrial disease (A)** In a *C. elegans dars-2*$^{-/-}$ model, there is no effect on mitochondrial stress, except for the highest dose of TAL. **(B)** Similarly to *dars-2*$^{-/-,}$ in a *C. elegans fars-2*$^{-/-}$ knockout none of the drugs had any effect at any concentration. **(C)** In a *C. elegans slc25A46(chp5)*$^{-/-}$ model, both the highest concentration of eltrombopag and of TAL rescued mitochondrial stress. **(D)** All three drugs show a trend towards rescue in fibroblasts from a patient with MT-ND1$^{-/-}$ but needs further replication before results can be assessed statistically. **(E)** In both healthy fibroblasts and fibroblasts from two different individuals with an ERCC6 mutation, patient 1 shows rescue at 0.1 µM OSI-906 and 1 µM TAL, while patient 2 shows rescue at 1 µM ELT. All panels show results of 24-hour treatment duration, with n=1 biological replicate/condition, and 3 technical replicates, unless otherwise specified. Cell viability was tested with a t-test with Benjamini-Hochberg for multiple test correction: *, $p < 0.05$; **, $p < 0.01$; ***, $p < 0.001$.

>IGFBP5_HUMAN
MVLLTAVLLLLAAYAGPAQSLGSFVHCEPCDEKALSMCPPSPLGCELVKEPGCGCCMTCA
LAEGQSCGVYTERCAQGLRCLPRQDEEKPLHALLHGRGVCLNEKSYREQVKIERDSREHE
EPTTSEMAEETYSPKIFRPKHTRISELKAEAVKKDRRKKLTQSKFVGGAENTAHPRIISA
PEMRQESEQGPCRRHMEASLQELKASPRMVPRAVYLPNCDRKGFYKRKQCKPSRGRKRGI
CWCVDKYGMKLPGMEYVDGDFQCHTFDSSNVE

**Figure S5. Sequence of the IGFBP5 mRNA** IGFBP5 LNP-mRNA was purchased from Perelman School of Medicine at the University of Pennsylvania. Length: 272, Mass (Da): 30,570.

**Supplemenntal Table Legends.**

**Table S1. SomaLogic Proteomics Differential Abundance.** Table of proteomics differential abundance results.

**Table S2. SomaLogic Proteomics Pathway Analysis.** Table of proteomics pathway analysis using WebGestaltR’s network-based topology analysis.
